# Top-down Proteomics of Myosin Light Chain Isoforms Define Chamber-Specific Expression in the Human Heart

**DOI:** 10.1101/2023.01.26.525767

**Authors:** Elizabeth F. Bayne, Kalina J. Rossler, Zachery R. Gregorich, Timothy J. Aballo, David S. Roberts, Emily A. Chapman, Wei Guo, J. Carter Ralphe, Timothy J. Kamp, Ying Ge

## Abstract

Myosin functions as the “molecular motor” of the sarcomere and generates the contractile force necessary for cardiac muscle contraction. Myosin light chains 1 and 2 (MLC-1 and -2) play important functional roles in regulating the structure of the hexameric myosin molecule. Each of these light chains has an ‘atrial’ and ‘ventricular’ isoform, so called because they are believed to exhibit chamber-restricted expression in the heart. However, recently the chamber-specific expression of MLC isoforms in the human heart has been questioned. Herein, we analyzed the expression of MLC-1 and -2 atrial and ventricular isoforms in each of the four cardiac chambers in adult non-failing donor hearts using top-down mass spectrometry (MS)-based proteomics. Strikingly, we detected an isoform thought to be ventricular, MLC-2v, in the atria and confirmed the protein sequence using tandem MS (MS/MS). For the first time, a putative deamidation post-translation modification (PTM) located on MLC-2v in atrial tissue was localized to amino acid N13. MLC-1v and MLC-2a were the only MLC isoforms exhibiting chamber-restricted expression patterns across all donor hearts. Importantly, our results unambiguously show that MLC-1v, not MLC-2v, is ventricle-specific in adult human hearts. Overall, top-down proteomics allowed us an unbiased analysis of MLC isoform expression throughout the human heart, uncovering previously unexpected isoform expression patterns and PTMs.

## Introduction

The hexameric protein complex myosin functions as the “molecular motor” of the sarcomere and generates contractile force through cross bridge formation with actin.^1, 2^ In cardiac tissue, myosin is composed of two heavy chains (MHC, either the α or β isoform) and four light chains (MLCs), which include two essential (ELC, or MLC-1) and two regulatory (RLC, or MLC-2) light chains.^3–5^ MLC-1 provides structural stability while MLC-2 regulates the function of the myosin motor through calcium binding and phosphorylation.^3^ MLC-1 and -2 each have two isoforms encoded by separate genes: *MYL4* and *MYL3* for MLC-1 and *MYL7* and *MYL2* for MLC-2.^6 7^

Each respective light chain has an ‘atrial’ and a ‘ventricular’ isoform so called because early studies suggested that the expression of these isoforms in the heart is chamber specific.^8, 9^ However, analysis of MLC expression in the heart was performed in animal models and, thus, the findings may not directly translate to humans.^10^ It is noteworthy that despite a lack of corroborating studies in humans, MLC-2v has become a commonly-used marker of ventricular specification in stem cell-derived cardiomyocyte (hPSC-CM) cultures.^11^ However, a direct investigation and confirmation of ventricular and atrial isoform expression in adult non-failing human hearts is lacking.

Top-down proteomics analyzes intact proteins, enabling unbiased analysis of proteoforms, a term describing all protein products produced from a single gene as a result of post-translational modifications (PTMs), sequence variants, and alternative splicing,^12^ as well as related proteins arising from different genes (e.g., isoforms not resulting from alternative splicing of a single gene).^12–17^ A direct analysis of MLC isoforms and proteoforms by top-down proteomics allows accurate quantification of their levels without *a priori* knowledge of sample contents. Previously, top-down proteomics was employed to characterize the amino acid sequences and N-terminal modifications of atrial and ventricular MLC isoforms from human and swine myocardial tissue.^18^

Herein, we employed top-down mass spectrometry (MS)-based proteomics to systematically evaluate differences in MLC proteoform and isoform expression in the four chambers of human donor hearts with no known history of cardiovascular disease (n=17). Surprisingly, our analysis revealed MLC-1v, but not MLC-2v, exhibited ventricle-specific expression, while MLC-2a exhibits atria-restricted expression. The exact amino acid sequence of MLC-2v in atrial tissue was then confirmed by online liquid chromatography-tandem MS (LC-MS/MS). Interestingly, we found a mixture of deamidated and non-deamidated MLC-2v species in the atria that was not observed in ventricular tissue. Overall, top-down proteomics provided us with an unbiased analysis of the MLC proteoform distribution in the four chambers of the non-failing adult human heart.

## Materials and Methods

### Human donor heart collection

Myocardial tissue from adult donor hearts with no history of cardiovascular disease were obtained from the University of Wisconsin Organ Procurement Organization (**Table S1** details clinical characteristics). Tissue collection procedures were approved by the Institutional Review Board of the University of Wisconsin-Madison (Study # 2013-1264). Cardiac tissue was dissected into left ventricle (LV), right ventricle (RV), left atrium (LA), and right atrium (RA) while fresh and immediately snap frozen in liquid nitrogen to preserve PTMs^19^ (**Figure S1**). The basal ventricular mid-wall was used as a representative region for LV and RV.

### Sarcomeric Protein Extraction

Sarcomeric proteins were extracted from human cardiac tissue using differential pH extraction as reported previously (**Figure S2**).^20, 21^ Total protein content was normalized by biochitonic assay and proteins were separated by online reversed phase liquid chromatography (LC) prior to MS analysis (LC-MS).

### Online LC-MS Analysis

LC-MS profiling was performed using a NanoAcquity ultra-high pressure LC system (Waters, Milford, MA) coupled to a high-resolution impact II quadrupole time-of-flight (Q-TOF) mass spectrometer (Bruker, Bremen, Germany).^19, 20, 22^ 400 ng total protein per injection were loaded on a home-packed PLRP column (250 μm i.d. x 200 mm length, PLRP-S, 10 μm particle size, 1000 Å pore size). Sarcomeric proteins were eluted using a flow rate of 4 μL/min over a gradient of 5% to 95% mobile phase B (0.1% formic acid in 50:50 acetonitrile and ethanol) over 40 minutes (mobile phase A: 0.1% formic acid in water). Electrospray ionization was used to introduce eluted proteins to the impact II Q-TOF mass spectrometer. Mass spectra were acquired at a scan rate of 0.5 Hz over 500-2000 *m/z* range. Injection replicates (n=3) were performed at the beginning of each run to ensure instrument reproducibility and extraction replicates (n=3) were analyzed to evaluate the reproducibility of sample preparation. Isotopic resolution was achieved for each analyte, and all proteins were identified with high mass accuracy (<3 ppm mass error).

Online tandem mass spectra were collected by data-dependent acquisition settings. The top 5 highest intensity precursors were selected and fragmented using collision-induced dissociation (CID). MS1 scan rate was 1 Hz and MS2 scan rate varied from 4-8 Hz depending on precursor ion intensity. Collisional energies were set to 18-45 eV. An active exclusion period of 1 min was used to maximize number of precursors selected. Electron Transfer Dissociation (ETD) was performed on the maXis II Q-TOF MS. The top 3 most abundant precursors in each MS1 spectrum were selected for fragmentation. The ETD settings were as follows: ion charge control ranging from 0-1000 ms, a reagent accumulation time of 25 ms, and an extended reaction time of 50 ms.

### Data Analysis

Mass spectra were analyzed using DataAnalysis v4.3 (Bruker Daltonics) and MASH Explorer.^23^ Mass spectra were averaged across retention time windows within the BPC to create MS1 spectra. Spectral deconvolution using the Maximum Entropy algorithm with a resolving power of 60,000 was performed to reveal the monoisotopic mass of the protein, and putative proteoform identifications are made based on accurate mass. Peaks whose molecular masses matched closely to predicted masses from the UniProtKB database were assigned the corresponding proteoform identity (**Table S3**). Proteoform quantitation is performed using peak intensities of unmodified and phosphorylated peaks, yielding a ratio of relative phosphorylation, P_total_ (mol P_i_/mol protein).^24 25^ Theoretical isotopic distributions of precursor and fragment ions were generated using IsoPro v3.0.

To better visualize isoforms over a wide retention time window, extracted ion chromatograms (EICs) were generated for quantitation (**Figure S3**). To quantitate relative intensities of isoforms in each sample, peak intensities from deconvoluted spectra were used if the isoforms eluted within the selected retention time window. To quantitate the relative intensity of isoforms in each sample, EICs were created from the top 5 most abundant ions (± 0.2 *m/z*) for each proteoform as described previously.^26^ Isoform intensity was normalized by dividing the peak areas under the curve (AUC) by the sum of peak areas for all isoforms pertaining to the protein family (e.g. MLC-1a/total MLC-1 content).^20^ For tandem MS sequence characterization, the monoisotopic mass of each fragment ion was analyzed against each putative protein sequence using IsoPro v3.0 using a mass error tolerance of 10 ppm. Isotopic distributions and theoretical fit were generated using IsoPro.

### Statistics

P_total_ values were analyzed for statistical significance using RStudio v1.2.5033. Differences between four chambers were evaluated using a linear model. One-way analysis of variance (ANOVA) was used to determine statistical significance between the four chambers. Tukey’s HSD *post hoc* test was used for a pairwise evaluation between regions and adjusted *p* values are reported. A 95% confidence interval was used for Tukey’s HSD. Differences between means were considered statistically significant if *p* <0.05. Levels of statistical significance are notated with an asterisk (*): \**p* ≤ 0.05, \*\**p* < 0.01, and \*\*\**p* < 0.001; no statistical significance *(ns)* if *p* > 0.05.

## Results

In this study, we employed top-down MS-based proteomics to comprehensively analyze the expression patterns of MLC-1 and -2 isoforms across the four chambers of non-failing human donor hearts (n=17) with no history of cardiovascular disease. Clinical characteristics of donors used in this study are presented in **Table S1**. Our platform efficiently captures the diversity of a complex protein mixture, enabling quantification of the relative abundance of proteoforms and isoforms in a single LC-MS experiment. This approach includes myocardial tissue homogenization, sarcomeric protein extraction, LC-MS/MS, and data analysis (**Figure S2**).

### Myofilament Protein Extraction and LC-MS/MS Platform is Highly Reproducible

Technical reproducibility of the online LC-MS/MS platform was demonstrated by alignment of retention times and intensity of base peak chromatograms (BPC) between injections (**Figure S4**). Method reproducibility was established by replicate extraction of proteins from the same cardiac chamber (n=3) (**Figure S5**). Proteoform quantitation was demonstrated by comparable intensity of the phosphorylated MLC-2v proteoform between extraction replicates (**Figure S5B**). Reproducible isoform quantitation was demonstrated by comparing the integrated area of MLC-2v EICs between technical replicates, ranging from 0.1-6% relative standard deviation across each cardiac chamber (**Figure S5C**). Finally, we demonstrated the highly reproducible detection of MLC-2v proteoform in atrial tissue despite the fact that the extraction and LC-MS sample runs were performed by different personnel (**Figure S6**). MS-based quantification of MLC expression was performed confidently after demonstration of high technical and extraction reproducibility.

### Top-down MS and MS/MS Analysis of MLC-1 Shows MLC-1v is Chamber-Specific

MLC-1a and MLC-1v were detected with high mass accuracy, matching their respective theoretical monoisotopic masses closely (**Figure 1A, Table S2**). MS analysis revealed single peaks corresponding to the ventricular and atrial MLC-1 isoforms in their respective chambers. The peak with a molecular mass of 21,828.91 Da detected in ventricular tissue matched well with the MLC-1v sequence in the UniProt/SwissProt database (Accession No. P08590-1, MYL3_HUMAN) when considering N-terminal methionine removal (−131.04 Da) and N^α^-trimethylation (+42.05 Da), consistent with previous reported N-terminal modifications (theoretical mass: 21,828.93 Da).^18^ The peak detected in atrial tissue with a molecular mass of 21,461.79 Da matched closely with the reported sequence of MLC-1a (Accession No. P12829-1, MYL4_HUMAN) when accounting for N-terminal methionine removal and N^α^-trimethylation as reported previously (theoretical mass: 21,461.87 Da).^18^

**Figure 1.**
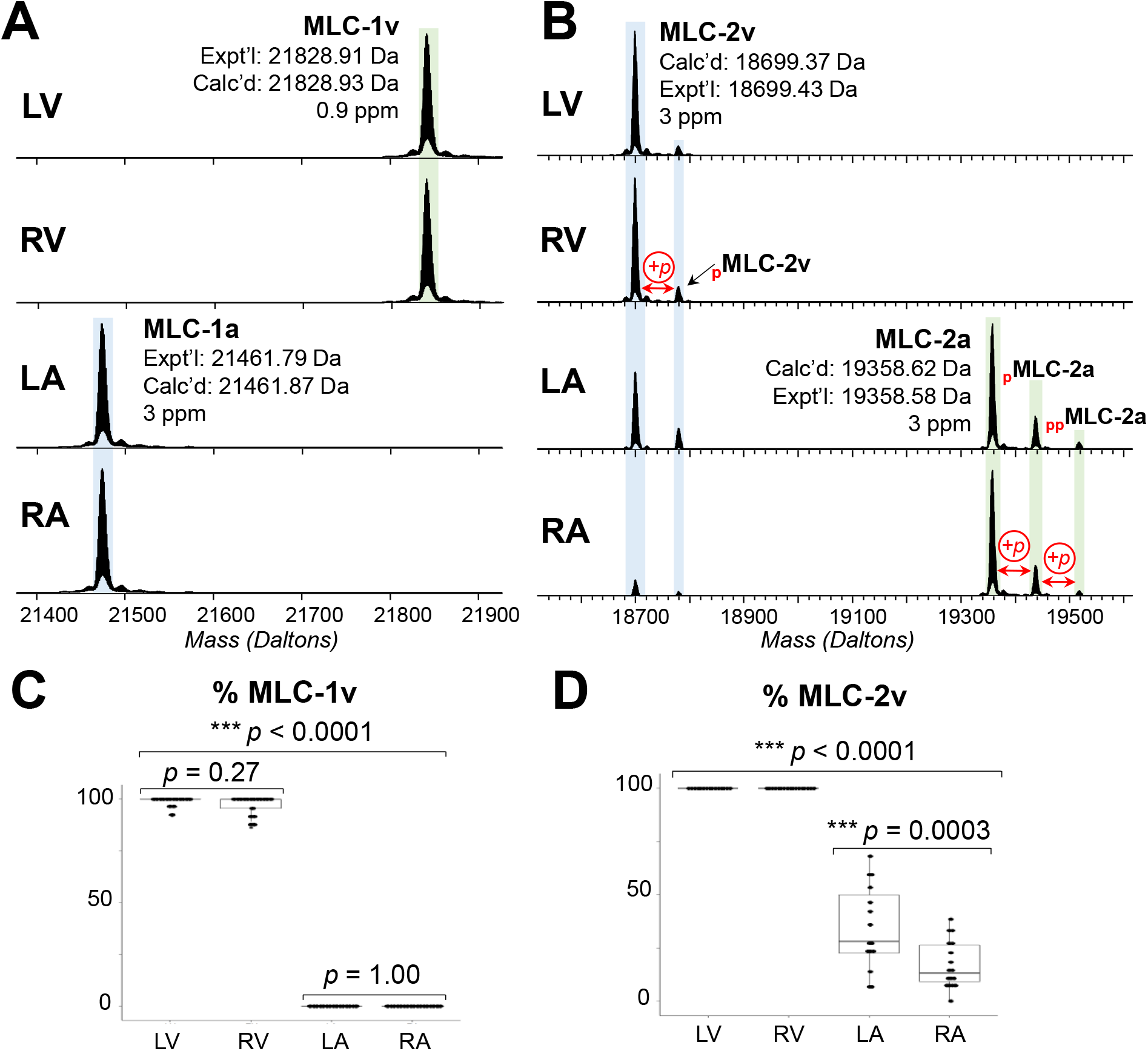
Top-down MS of MLCs Shows MLC-1v is Ventricle-specific. **A)** Deconvoluted spectra showing expression of ventricular and atrial isoforms of myosin light chain 1 (MLC-1). MLC-1v is the dominant isoform in LV and RV tissues and was exclusively detected in LV and RV. MLC-1a was the only isoform detected in LA and RA tissues. **B)** Deconvoluted spectra showing ventricular and atrial isoforms of myosin light chain 2 (MLC-2). MLC-2v was consistently detected throughout all four chambers of the human heart. MLC-2v was the dominant isoform in LV and RV and detected at varying levels in LA and RA. The atrial isoform MLC-2a was detected exclusively in LA and RA. Isotopic resolution was achieved for each of the isoforms and all proteins were identified with high mass accuracy (3 ppm mass error). **C)** Quantitation of the expression of MLC-1v relative to total MLC-1a+MLC-1v in LV, RV, LA, and RA (n=17 donors). Statistically-significant differences across the four chambers were determined by one-way ANOVA followed by Tukey’s HSD *post hoc* test for pairwise comparisons (p ≤ 0.05). One-way ANOVA resulted in overall *p* value of 2.2e-16 (**Table S3**). Results of pairwise comparisons are reported on the plot and are notated with an asterisk (*): \**p* ≤ 0.05, \*\**p* < 0.01, and \*\*\**p* < 0.001; no statistical significance if *p* > 0.05. **D)** Quantitation of the expression of MLC-2v relative to total MLC-2a+MLC-2V in LV, RV, LA, and RA (n=17 donors). Statistically-significant differences across the four chambers were determined by one-way ANOVA followed by Tukey’s HSD *post hoc* test for pairwise comparisons (p ≤ 0.05). One-way ANOVA resulted in overall *p* value of 2.0e-16 (**Table S3**). Results of pairwise comparisons are reported on the plot and are notated with an asterisk (*): \**p* ≤ 0.05, \*\**p* < 0.01, and \*\*\**p* < 0.001; no statistical significance if *p* > 0.05.

LC-MS/MS generated N- and C-terminal protein fragment ions using CID. Fragment ions confirmed the MLC-1 isoform sequences with N-terminal modifications as previously reported.^18^ Ions corresponding to MLC-1v, including *b_23_* and *y*_80_, were observed consistently across LV and RV preparations from different hearts (**Figure S7A**), matching closely to theoretical isotopic distributions with high mass accuracy. In total, 22 *b* and 16 *y* ions were identified within ≤ 10 ppm mass error, corresponding to 87% sequence coverage of MLC-1v from a single LC-MS/MS run (**Figure S8**). Similarly, key N- and C-terminal fragments matching MLC-1a were identified with consistency across LA and RA preparations from different hearts, matching theoretical isotopic distributions with high mass accuracy (**Figure S7B**). MLC-1a fragmentation by CID yielded 31 total *b* and 15 *y* ions within ≤ 10 ppm mass error (**Figure S9**).

Quantification of MLC-1 isoforms revealed MLC-1v was ventricle-specific. MLC-1v was the most abundant isoform in LV and RV, averaging 99 ± 0.6% MLC-1v in LV and 97 ± 1.1% in RV (**Figure 1C**). Close analysis of the raw mass spectra showed MLC-1v was not detected above baseline noise in LA and RA (**Figure S10**). One-way ANOVA was used to evaluate MLC-1v expression across the four chambers (*p* = 2.2e-16), and pairwise comparisons revealed statistically significant contrasts between LA and RA when compared to LV and RV (*p* < 0.0001 for each comparison, for details refer to **Table S3**). No lateral distinctions were noted in the ventricles and atria (LV-RV *p* = 0.27; LA-RA *p* = 1.00).

### Ventricular Isoform of MLC-2 Detected in Atrial Tissues Using LC-MS/MS

Top-down MS revealed 2 proteoforms related to MLC-2v and 3 proteoforms related to MLC-2a. Two proteoforms with molecular masses of 18,699.40 and 18,768.36 Da were detected in each cardiac chamber (**Figure 1B, Table S2**). The peak with a mass of 18,699.40 Da matched well to the calculated molecular mass of MLC-2v (Accession No. P10916-1, MLRV_HUMAN) after considering N-terminal methionine cleavage and N^α^-trimethylation as previously reported. The peak of 18,768.36 Da was an additional +79.97 Da from MLC-2v, corresponding to the addition of meta-phosphoric acid HPO_3_ and, thus, was assigned as mono-phosphorylated *p*MLC-2v. The remaining three proteoforms were exclusively detected in atria and had molecular masses of 19,346.54, 19,426.50, and 19,506.47 Da. The 19,346.54 Da peak matches with predicted molecular mass of MLC-2a (Accession No. Q01449-1, MLRA_HUMAN) after considering N-terminal methionine cleavage and N^α^-acetylation (+42.01 Da). Mono-phosphorylated *_p_*MLC-2a was assigned to the peak of 19,426.50 Da due to an addition of +79.97 Da from MLC-2a and bis-phosphorylated *_p_*MLC-2a was assigned to peak of 19,506.47 Da due to a mass difference of +159.91 Da. Each isoform showed good fitting of theoretical isotopic abundance and distribution of the isotopomer peaks (**Figure S12**), including phosphorylated proteoforms (**Figure S13**).

MS/MS enabled by CID was used to confirm the identity of MLC-2v across the four cardiac chambers. A total of 47 MLC-2v fragment ions were identified across LV, RV, LA, and RA, including 22 *b* and 25 *y* ions (**Figure S14**). For example, the *y_24_* ion corresponding to C-terminal MLC-2v matched closely to the theoretical isotopic distribution for each cardiac chamber, and the experimentally-determined mass matches the theoretical mass with sub-ppm accuracy, indicating agreement with previously reported MLC-2v sequence (**Figure S15A**). Due to considerable sequence heterogeneity between MLC-2v and MLC-2a,^27^ the observed fragment ions overwhelmingly indicate the presence of MLC-2v in the atrial chambers and do not match with a truncated or alternatively-spliced MLC-2a isoform. LC-MS/MS revealed key fragment ions matching to the reported sequence of MLC-2a, such as the *y_25_* ion shown across left and right atrial tissues (**Figure S15B**). In sum, 37 *b* and 12 *y* ions were matched to the MLC-2a sequence within ≤ 10 ppm mass error, resulting in 61% sequence coverage (**Figure S16**).

Quantification of MLC-2 isoforms revealed significant differences of MLC-2v expression across the four cardiac chambers. In LV and RV, MLC-2v remained at 100% of total signal. MLC-2v content ranged widely in atrial tissues, from 6-68% in LA and 0-39% in RA, averaging 34 ± 5.1% MLC-1v in LA and 17 ± 2.7% in RA. MLC-2a was not detected above baseline noise in LV and RV, as shown by close examination of each raw mass spectrum (**Figure S12**). MLC-2v expression across the four chambers was significant by one-way ANOVA (*p* = 2.0e-16) (**Figure 1D**). Pairwise comparisons between LV, RV, LA, and RA revealed significant differences between ventricles and atria (all pairwise comparisons *p* < 0.0001 by Tukey HSD). The % MLC-2v was also statistically significant laterally, between LA and RA (*p* = 0.0003).

No chamber-related differences in relative phosphorylation (mol P_i_/mol protein, or P_total_) of MLC-2 isoforms were detected. The relative phosphorylation of MLC-2v averaged 0.15 ± 0.02 P_total_ in LV and 0.13 ± 0.02 P_total_ in RV, and 0.19 ± 0.03 P_total_ in LA and 0.16 ± 0.03 P_total_ in RA. These values did not yield a significant difference across the four chambers (*p* = 0.48) when compared by one-way ANOVA (**Figure S13C**). Similarly, a comparison of the relative phosphorylation of MLC-2a between the left and right atria (avg: 0.25 ± 0.03 P_total_ in LA and 0.22 ± 0.03 P_total_ in RA) was not significantly different (*p* = 0.52 by Student *t* test).

### Atrial-Specific Deamidation of MLC-2v Localized to N13 by MS/MS

Top-down MS-based proteomics identified and localized an atrial-specific deamidation of human MLC-2v. A difference of +0.98 Da was found in the measured monoisotopic masses of MLC-2v in atrial tissues compared to ventricular tissue (**Figure 2A**). N-terminal fragment ions produced by CID MS/MS revealed a +0.98 Da mass shift near the N-terminus of the protein. This phenomenon was observed in all N-terminal fragments produced by CID MS/MS in atrial tissue, including the shortest fragment *b*_15_ (**Figure 2B**). Putative PTMs matching the observed mass shift are deamidation of asparagine^28^ or citrullination of arginine.^29^ Since both asparagine and arginine residues are in the first 15 aa residues of MLC-2v sequence, fragmentation closer to the N-terminus was needed to localize and identify the PTM. However, the CID spectra contained no N-terminal fragments shorter than *b*_15_, possibly due to the basic nature of amino acid residues, which may have inhibited fragmentation by CID by conferring resonance-induced stability to the peptide backbone.^30^

**Figure 2.**
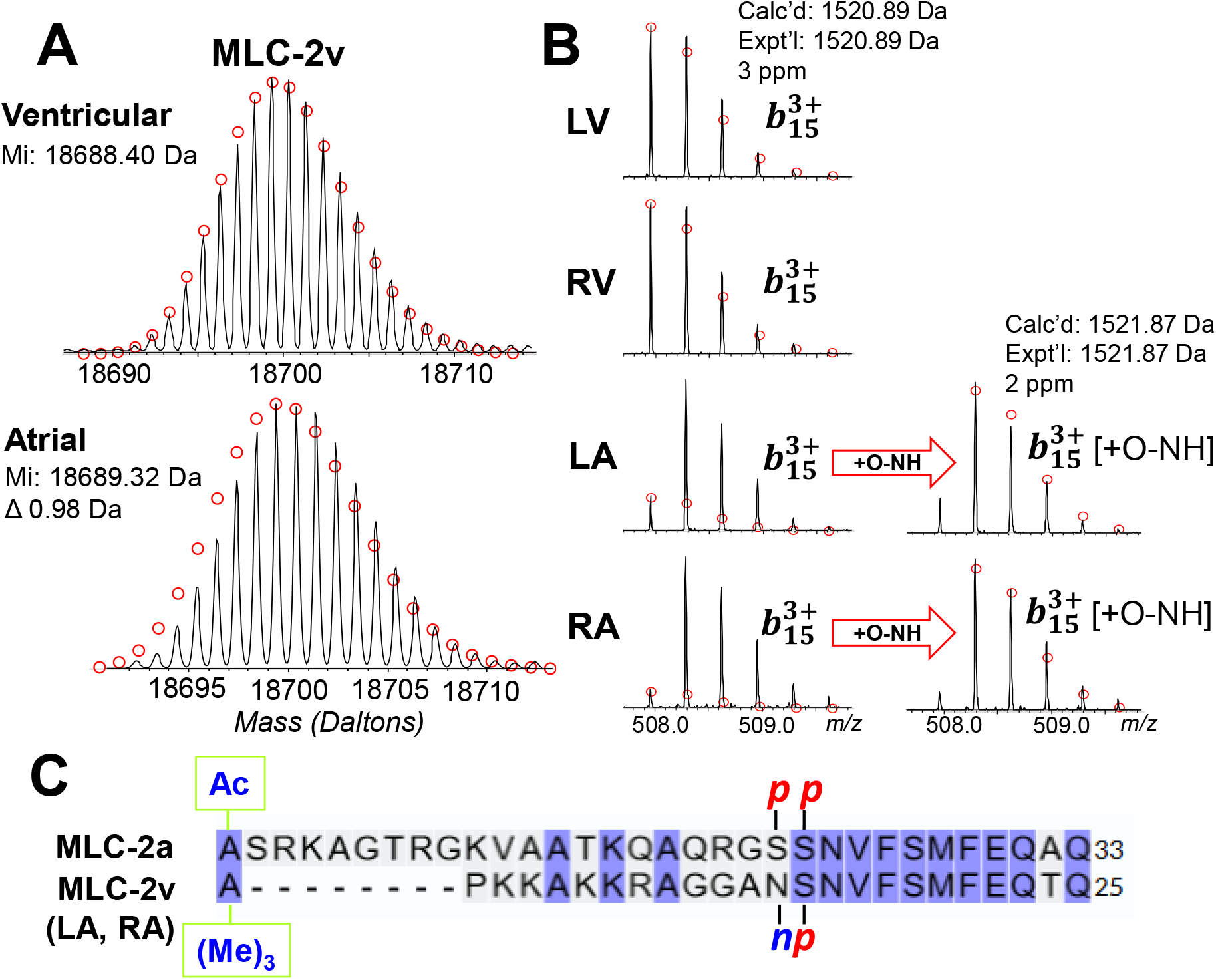
Atrial-Specific Deamidation of MLC-2v Localized to N13 by MS/MS. **A)** Deconvoluted spectra of MLC-2v showing a difference of 0.98 Da in monoisotopic mass detected between ventricular and atrial tissues. **B)** Fragment ion spectra of *b*_15_, the smallest N-terminal fragment of MLC-2v observed by CID MS/MS. Deamidation (+0.984 Da) was added to the theoretical mass of ion *b*_15_ and the resulting theoretical isotopic distribution was fitted in LA and RA. **C)** Sequence alignment of the first 33 residues of MLC-2a (above) and 25 residues of MLC-2v (below) with deamidation indicated at the N13. Sequence alignment shows site of phosphorylation at Ser22 in MLC-2a matches with site of deamidation at N13 in MLC-2v sequence.

To further localize the putative modification, we utilized ETD to generate shorter N-terminal fragments. ETD revealed shorter N-terminal fragments including *c*_7_, *c*_8_, *c*_9_, and *c*_12_ that matched well to calculated isotopic distribution of MLC-2v sequence and displayed comparable intensities to the same fragment ions observed in LV and RV (**Figure S17A**). This result eliminates the possibility of citrullination at the Arg8 residue, indicating the PTM is located between residues aa[12-15]. A *c*_14_ ion showed a reduction of monoisotopic intensity and aligned more closely to the MLC-2v aa[1-14] when accounting for deamidation [+O-NH] (**Figure S17B**), indicating the PTM was located between aa[12-14]. We assigned the putative PTM corresponding to +0.98 Da as deamidation located at Asp13 based on the identity of isolated residues and mass shift. We conclude there is a mixture of deamidated and unmodified MLC-2v proteoform species in the atrial tissues due to the observed overlapping isotopic envelopes of N-terminal fragment ions. Since this PTM was only detected in LA and RA (LV or RV had normal isotopomer distribution matching the predicted N-terminal sequence), the observed atrial deamidation is not an artefactual or preparation-based modification, but one with potential biological relevance.

## Discussion

In this study, we characterized the spatial distribution of MLC isoforms across ventricular and atrial tissue of adult non-failing human hearts. To identify and quantify the MLC isoforms in different chambers from human cardiac tissue, we used top-down MS-based proteomics to enable a “bird’s eye view” of the intact MLCs, revealing unique patterns of isoform expression and PTMs across the four chambers of the human heart.

### MLC-1v, But Not MLC-2v, Is Ventricle-Specific in Human Hearts

In the mature heart, MLC atrial and ventricular isoforms are widely considered to be chamber-specific,^6, 7^ but in this study top-down MS-based proteomics has uncovered evidence of expression of MLC-2v in human atrial tissues unambiguously. **Figure 1** shows the detection and quantification of MLC isoforms across LV, RV, LA, and RA. We found MLC-1v but not MLC-2v exhibits ventricular-restricted expression. Peaks matching molecular masses of MLC-2v and *p*MLC-2v were identified in atrial tissue. Strikingly, MLC-2v was detected in the atria of every heart from all 17 donors with no cardiac diseases. The %MLC-2v of total MLC-2 content was quantitated and deemed statistically significant by one-way ANOVA, with significant differences between ventricular and atrial samples and a lateral difference between LA and RA.

MLC-1a and -1v have well-established functional differences in both actin-myosin tethering and cross-bridge cycling kinetics.^3^ Ventricular expression of MLC-1a has been found under states of stress such as cardiomyopathy.^31^ It is possible MLC-2v functions as a similar adaptive isoform switch during a state of cardiac stress similarly to MLC-1. MLC-2v has been detected in atrial tissue in previous studies of pressure overload in both humans^32^ and rats.^33^ Interestingly, MLC-2v was also found in atria of adult spontaneously hypertensive rats without induced pressure overload, suggesting the expression of the ventricular isoform in atrial tissue was the result of a predetermined phenotype rather than a direct response to pressure overload.^33^ Functionally, ventricular and atrial isoforms of MLC-2 have been shown to affect the morphogenesis and cross bridge cycling of the myocardium. In transgenic mice, replacement of MLC-2a by -2v in atria has shown enhanced contractile generation and calcium sensitivity similar to the ventricular cardiomyocyte controls.^34, 35^ A nuanced assessment of specific isoform influence on either atrial or ventricular function, and the regulation of their differential expression in healthy and diseased states, remains to be explored.

### Identification of Atrial-specific MLC-2v Deamidation on Asn13

We detected a small mass difference of <1 Da in MLC-2v in the atrial tissues compared to the ventricles. **Figure 2** details the localization and identification of this PTM using a combination of fragmentation methods (CID and ETD). Deamidation of MLC-2v from non-failing human cardiac tissue has been previously reported by analysis of enriched tryptic peptides by tandem MS.^36^ For the first time, we have localized deamidation to the Asn13 residue in human MLC-2v using a combination of ETD and CID. This site is consistent with a previous assignment of Asn13 as the site of deamidation of MLC-2v in rabbit myocardium^37^ and in non-human primate skeletal muscle (MLC-2s isoform).^38^

The site of deamidation is adjacent to the regulatory phosphorylation site Ser14. The additional negative charge imparted by deamidation^39^ adjacent to the site of phosphorylation mimics the pattern of bis-phosphorylation of MLC-2a at sites Ser22/23 (**Figure 2C**). Atrial-specific deamidation of MLC-2v may have functional implications relating to cardiac force generation in the atrial chambers. Phosphorylation induces a conformational change in MLC-2,^36^ and the addition of a second negative charge may serve to heighten calcium sensitivity and induce further conformational change to the protein. Further investigation to determine the functional consequences of atrial-specific deamidation in human MLC-2v may help elucidate the role of partial replacement from MLC-2a to MLC-2v in the atria.

### Implications for ‘Chamber-specific’ *in vitro* hPSC-CM Models

The assumptions regarding the specificity of MLC isoform expression and distribution is increasingly exploited as a marker for either atrial or ventricular cardiomyocyte differentiation from cultured human pluripotent stem cells (hPSCs). As hPSC-CM maturation strategies advance,^40^ there is a growing need to understand sarcomeric isoform distribution in healthy adult human hearts to create accurate models. For example, atrial fibrillation is a common cardiac arrhythmia in adults leading to serious health complications,^41^ and atrial-exclusive populations of hPSC-CMs have been used to model its pathophysiology and effects on function.^42 43^ While the absence of MLC-2v expression is often used to verify atrial-like hPSC-CMs,^42^ the present results indicate MLC-2v can be found alongside MLC-2a in human adult atrial tissue. For applications seeking to closely resemble adult atrial cardiomyocyte physiology, the presence, or absence, of MLC-2v may be an important consideration since there are known isoform-specific effects on contractile performance form both animal and human disease models.^34, 44^

It is important to note that MLC-2v is a commonly-used marker of ventricular specification in hPSC-CM cultures, with the assumption that its absence is indicative of successful atrial cardiomyocyte differentiation.^11,45^ The data presented here suggest the absence of MLC-1v may be a more accurate indicator than MLC-2v to confirm atrial cardiomyocytes, as MLC-1v was found exclusively expressed in human ventricular tissue. MLC-1v has not been used as a marker for hPSC-CM differentiation and maturation in the stem cell community to date.

## Conclusion

In this study, we characterized the chamber-specific distribution of MLC isoforms in adult non-failing human hearts (n=17) using top-down MS-based proteomics. MLC atrial and ventricular isoforms are widely considered to be chamber-specific,^6, 7^ but top-down MS-based proteomics has uncovered evidence of expression of MLC-2v in human atrial tissues. Top-down proteomics enables an unbiased analysis of isoform distribution in the human heart by directly analyzing intact proteins to reveal PTMs. Here, we found MLC-1v but not MLC-2v exhibits ventricular-restricted expression. We confirmed the protein sequence of MLC-2v in the atria using high resolution top-down MS/MS and detected a deamidated species of MLC-2v exclusive to atrial tissue. The modification was localized to the N13 residue in MLC-2v. Such “bird’s eye view” of the proteome by top-down proteomics can unveil previously unattainable knowledge of isoform distribution in the heart.

## Supporting information

EIC Quantification Data

Ptotal Quantification Data

## Data Availability

Raw data are available in the MassIVE repository with identifier MSV000091143.

## Acknowledgements

We are grateful to the donors and their families for their generous donation of tissue. We thank those who assisted with cardiac tissue collection and dissection, especially Tim Zhou, MD. We thank James Anderson and Carrie Sparks at the University of Wisconsin Organ and Tissue Donation for coordination of donor heart collection. Financial support was provided by National Institutes of Health (NIH) R01 HL096971 (Y.G.) and R01 HL109810 (Y.G.). Y.G. also acknowledges R01 GM117058, R01 GM125085, and S10 OD018475. E.F.B. acknowledges support from R01 HL148059-01. K.J.R acknowledges the National Science Foundation Graduate Research Fellowship Program under Grant No. DGE-1747503 and the Graduate School and the Office of the Vice Chancellor for Research and Graduate Education at the University of Wisconsin-Madison, funded by Wisconsin Alumni Research Foundation. D.S.R. acknowledges support from the American Heart Association Predoctoral Fellowship Grant No. 832615/David S. Roberts/2021. T.J.A. acknowledges support from the NIH Training Grant T32 GM008688. E.A.C. acknowledges support from the NIH Chemistry-Biology Interface Training Program NIH T32GM008505. W.G. and Z.G. acknowledge funding from the NIH NHBLI HL148733 and the American Heart Association Foundation 19TPA3480072. J.C.R. acknowledges support from R01 HL139883. Any opinions, findings, and conclusions or recommendations expressed in this material are those of the authors and do not necessarily reflect the views of the National Science Foundation.

