## Supplemental Information for "Top-down Proteomics of Myosin Light Chain Isoforms Define Chamber-Specific Expression in the Human Heart"

### Table of Contents

#### Supplemental Tables


- Table S1. Donor clinical characteristics of n=17 hearts provided by the UW Hospital Organ Procurement Organization.
- Table S2: Myosin light chain proteoforms identified from human cardiac tissue.
- Table S3:  Results of Statistical Analysis.


#### Supplemental Figures


- Figure S1. Dissection of a human donor heart.
- Figure S2. Quantitative Analysis of Sarcomeric Proteins using Top-down MS.
- Figure S3. Quantitation of MLC-1 using Extracted Ion Chromatograms (EICs).
- Figure S4. Injection Reproducibility of LC-MS platform.
- Figure S5. Reproducible LC-MS analysis of sarcomeric proteins from multiple regions in the human heart.
- Figure S6. MLC-2v detected in LA tissue across multiple extractions, sample runs, and analysts.
- Figure S7. MLC-1 Fragment Ions identified in ventricular and atrial tissue.
- Figure S8. Online LC-MS/MS of MLC-1v. s
- Figure S9. Online LC-MS/MS of MLC-1a.
- Figure S10. Raw mass spectra of isoforms of MLC-1 in LV, RV, LA, and RA.
- Figure S11. Figure S8. MLC-1a detected at low levels in some ventricular donor tissues.
- Figure S12. Raw mass spectra of isoforms of MLC-2 in LV, RV, LA, and RA.
- Figure S13. Identification and quantitation of phosphorylated proteoforms of MLC-2 across the four cardiac chambers.
- Figure S14. Online LC-MS/MS of MLC-2v.
- Figure S15. Online LC-MS/MS of MLC-2a.
- Figure S16. MLC-2 fragment ions identified in ventricular and atrial tissue.

**Table S1. Donor clinical characteristics of n=17 hearts provided by the UW Hospital Organ Procurement Organization.** All donors had no history of cardiac disease and were not used for transplant due to antigen incompatibility, transplant candidate availability, or co-morbidity. Average and Standard Error (SEM) is reported for age. Number of donors (n) is indicated in parentheses after each reported percentage.

| Clinical Characteristic | Percentage of sample cohort |
|---|---|
| Age | 46 ± 0.7 years<br>min: 24 years<br>max: 64 years |
| Male<br>Female | 47% (n = 8)<br>53% (n = 9) |
| Hypertension | 47% (n = 8) |
| Diabetes | 12% (n = 2) |
| Hyperlipidemia | 18% (n = 3) |
| Stroke | 18% (n = 3) |
| Substance Abuse | 24% (n = 4) |
| Smoking | 24% (n = 4) |

**Table S2: Myosin Light Chain Proteoforms Identified from Human Cardiac Tissue**. Proteoform name, gene, Uniprot ID, modifications, calculated (Calc'd) and experimental (Expt'l) monoisotopic masses, and error in parts per million (ppm) are reported. Abbreviations: myosin light chains 1 (MLC-1) and 2 (MLC-2). Modifications: N-terminal methionine removal (-Met), acetylation (+Ac), tri-methylation $(+Me)_3$, mono-phosphorylation (+P), and bis-phosphorylation (+2P).

| Proteoform | Gene | Uniprot ID | Modifications | Calc'd (Da) | Expt'l (Da) | Error (ppm) |
|---|---|---|---|---|---|---|
| MLC-1v | MYL3 | P08590-1 | $+(Me)_3$-Met | 21828.93 | 21828.91 | -0.9 |
| MLC-1a | MYL4 | P12829-1 | $+(Me)_3$-Met | 21461.87 | 21461.79 | -3.3 |
| MLC-2a | MYL7 | Q01449-1 | +Ac-Met | 19346.59 | 19346.54 | -3.0 |
| pMLC-2a | | | +P, +Ac-Met | 19426.56 | 19426.50 | -3.0 |
| ppMLC-2a | | | +2P, +Ac -Met | 19506.53 | 19506.47 | -2.9 |
| MLC-2v | MYL2 | P10916-1 | $+(Me)_3$ -Met | 18688.38 | 18688.40 | 1.1 |
| pMLC-2v | | | +P, $+(Me)_3$ -Met | 18768.35 | 18768.36 | 0.8 |

**Table S3: Results of Statistical Analysis**. One-way analysis of variance (ANOVA) was used to determine statistical significance between cardiac regions. Tukey's HSD *post hoc* test was used for a pairwise evaluation between regions and adjusted *p* values are reported. A 95% confidence interval was used for Tukey's HSD. Levels of statistical significance are notated with an asterisk (*): *$p \leq 0.05$, **$p < 0.01$, and ***$p < 0.001$; no statistical significance *(ns)* if $p > 0.05$.

| | **Mean** | **ANOVA F-value** | ***p*-value** | **Contrasts where $p \leq 0.05$** | **Contrasts where $p > 0.05$** |
|---|---|---|---|---|---|
| **% MLC-1v** | LV: 92.3%<br>RV: 86.3%<br>LA: 0%<br>RA: 0% | 7448.8 | < 2.2e-16 *** | LA-LV***, LA-RV***, LV-RA***, RA-RV*** | LA-RA, LV-RV |
| **% MLC-2v** | LV: 100%<br>RV: 100%<br>LA: 34.4%<br>RA: 17.2% | 247.5 | <2.0e-16 *** | LA-LV***, LA-RA***, LA-RV***, LV-RA***, RA-RV*** | LV-RV |
| **$P_{total}$ MLC-2v** | LV: 0.15<br>RV: 0.13<br>LA: 0.19<br>RA: 0.16 | 0.8356 | 0.48 | N/A | N/A |
| **$P_{total}$ MLC-2a** | LA: 0.25<br>RA: 0.22 | 0.4292 | 0.52 | N/A | N/A |

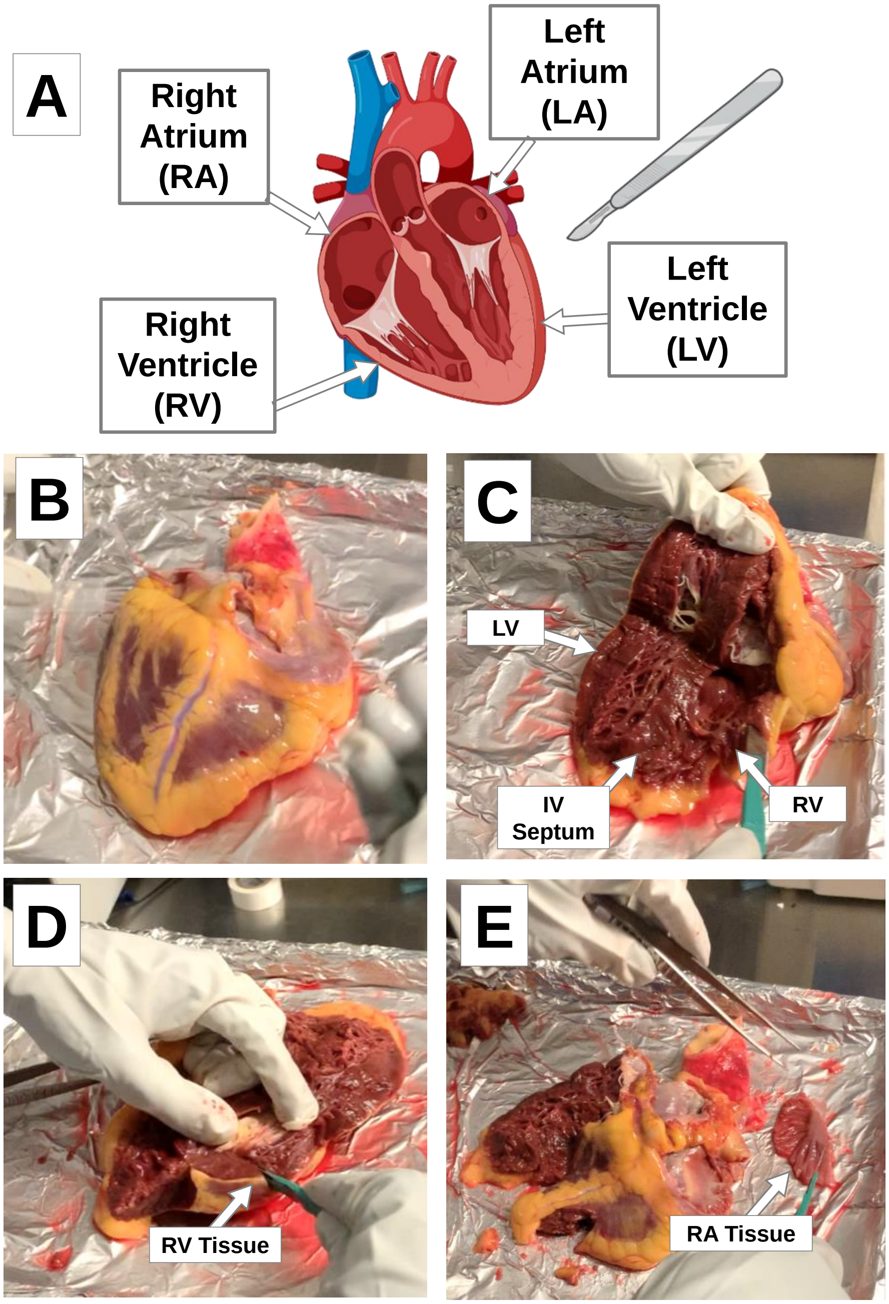


**Figure S1. Dissection of a human donor heart**. **A)** Regions dissected from myocardial tissue included the left ventricle (LV), right ventricle (RV), left atrium (LA), and right atrium (RA). **B)** Photograph of human donor heart prior to dissection. Donor hearts are stored in cardioplegic solution and packed in ice during transfer from organ recovery surgical suite to research lab. The heart is placed on a metal plate on top of a container of dry ice and the dissection is performed at 4 ºC. **C)** Initial cut of the dissection is performed by slicing open the heart from apex. The Interventricular Septum (IV Septum) separates the left and right ventricles, and the free walls are designated as LV and RV. **D)** LV and RV tissue is carefully excised and flash frozen in liquid nitrogen. **(E)** Lastly, the LA and RA are removed and flash frozen in liquid nitrogen.

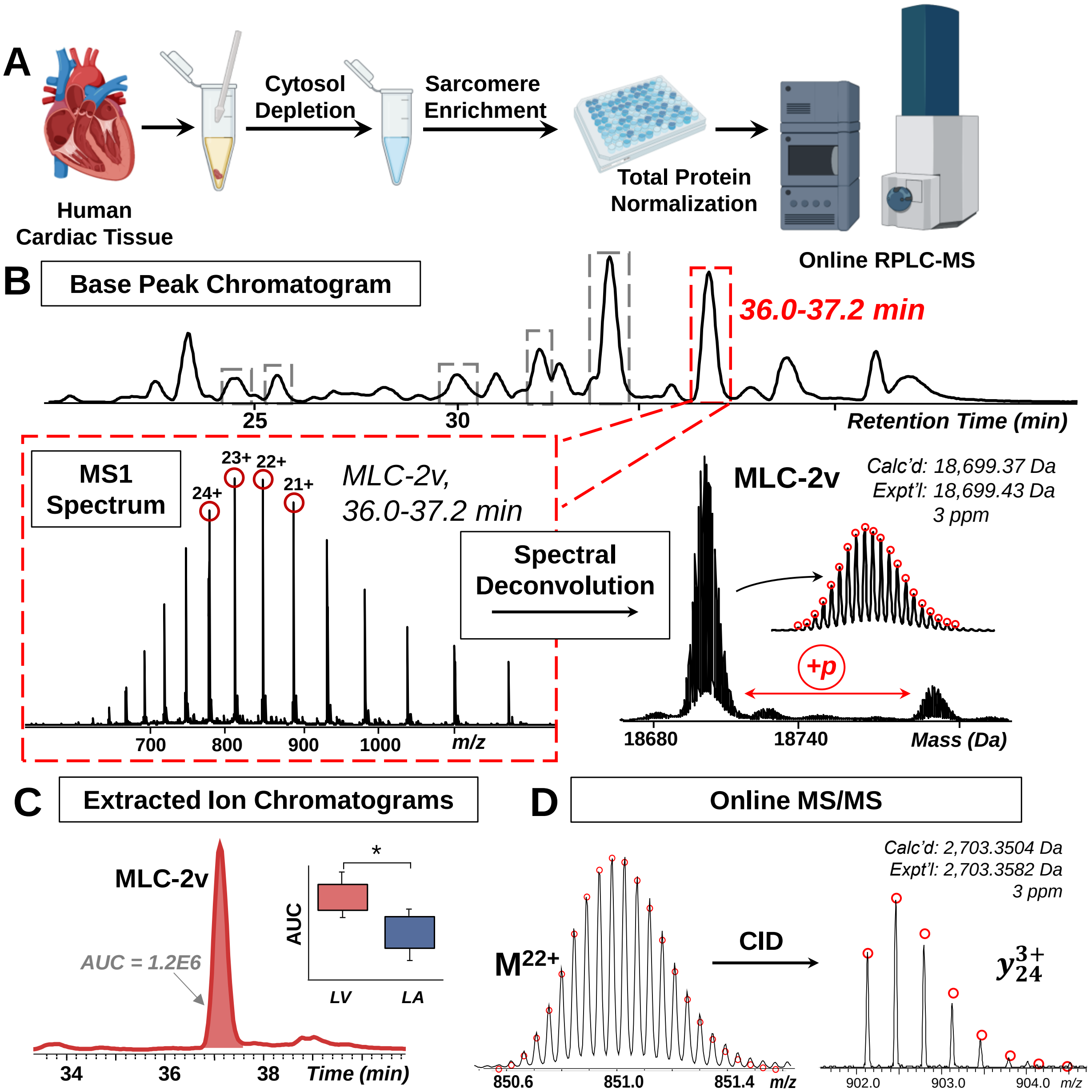


**Figure S2. Quantitative Analysis of Sarcomeric Proteins using Top-down Mass Spectrometry. A)** Workflow of the analysis of sarcomeric proteoforms from human cardiac tissue. This approach includes myocardial tissue homogenization, a differential pH-based enrichment for the sarcomeric proteome, total protein normalization, and online RPLC-MS. **B)** Representative RPLC-MS analysis of sarcomeric proteins. Base peak chromatogram (BPC) demonstrates efficient separation of intact proteins from the complex mixture. MS spectra for MLC-2v are averaged across selected retention time windows (highlighted in red), ranging from 36.0-37.2 min. The compiled MS1 spectrum shows the protein charge state distribution. Spectral deconvolution is performed to reveal the monoisotopic mass of the protein. **C)** To quantitate the relative abundance of MLC isoforms across a large retention time window, Extracted Ion Chromatograms (EICs) are created using the top 5 most abundant ions corresponding to each proteoform, and the area under the curve (AUC) is integrated and used for quantitation. **D)** Tandem MS **(**MS/MS) is performed to confirm proteoform identity. Precursor ions (charge state $M^{22+}$ shown) are selected for fragmentation using Collisionally Induced Dissociation (CID), generating N- and C-terminal fragments matching to a theoretical fragment mass and isotopic distribution.

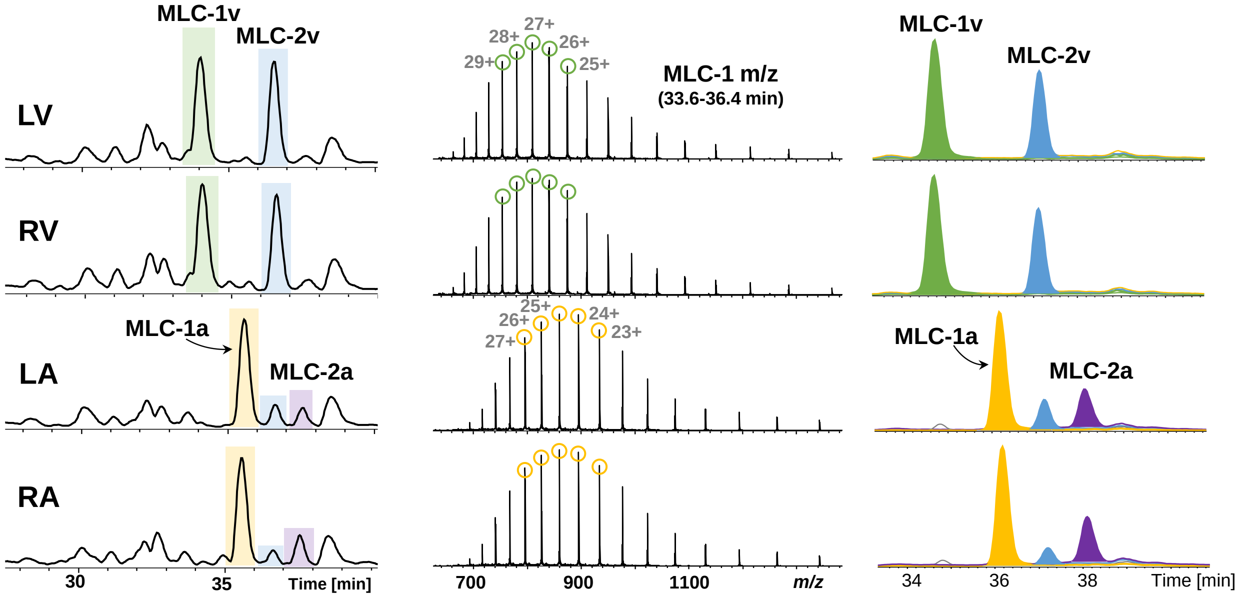


**Figure S3. Quantitation of MLC-1 using Extracted Ion Chromatograms (EICs). A)** Stacked overlay of BPCs from LV, RV, LA, and RA tissue. **B)** Averaged mass spectrum showing MLC-1. The top 5 most intense charge states for MLC-1v (green) and MLC-1a (yellow) are selected (average ± 0.2 *m/z*) and then extracted as a single peak corresponding to each isoform. **C)** EICs for MLC-1 and MLC-2 isoforms. The AUC for each peak was integrated and used for quantitation.

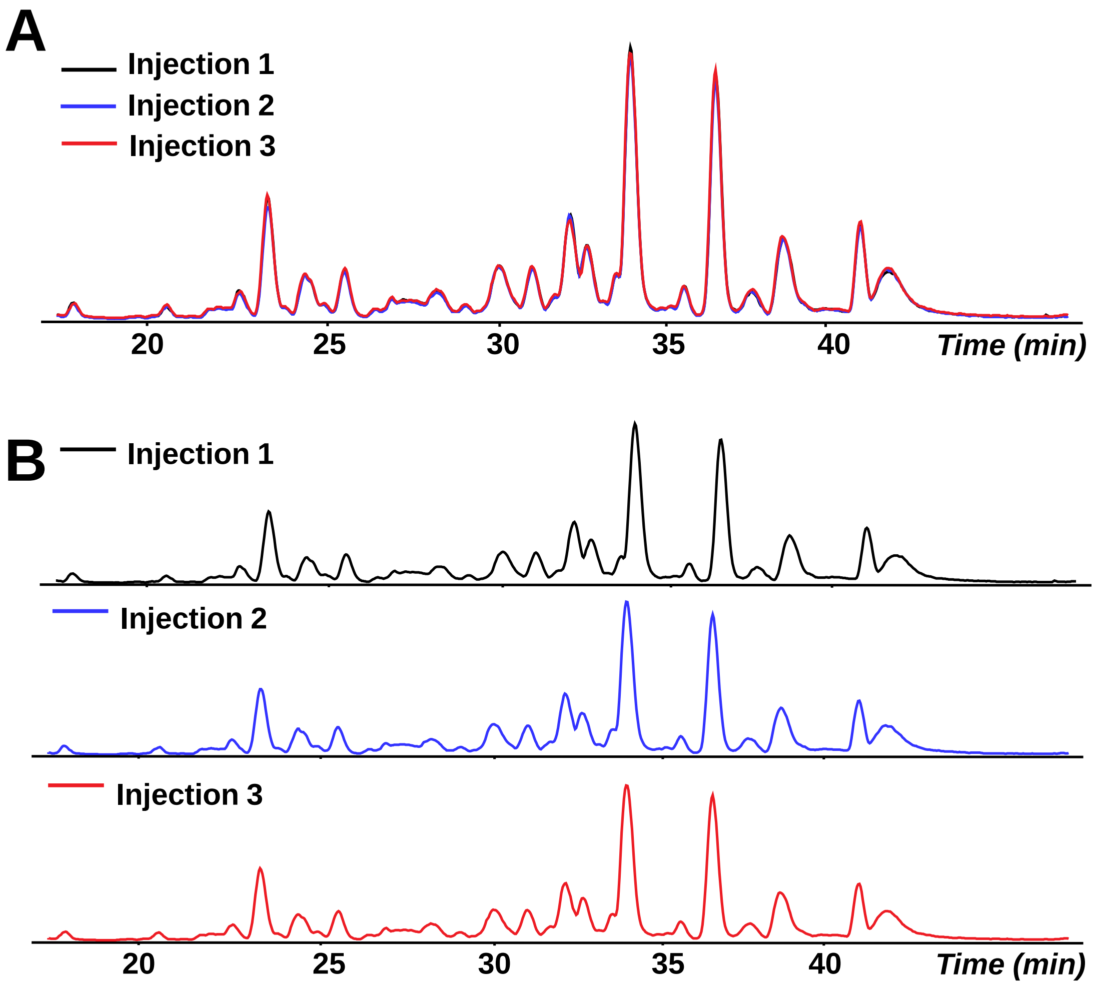


**Figure S4**. **Injection Reproducibility of Liquid Chromatography (LC)-MS platform**. **A)** Overlay and **B)** stacked overlay of injection replicates of sarcomere extract enriched from LV. Retention times and intensity of base peak chromatograms are nearly identical between injection replicates, demonstrating high technical reproducibility of the online RPLC-MS platform.

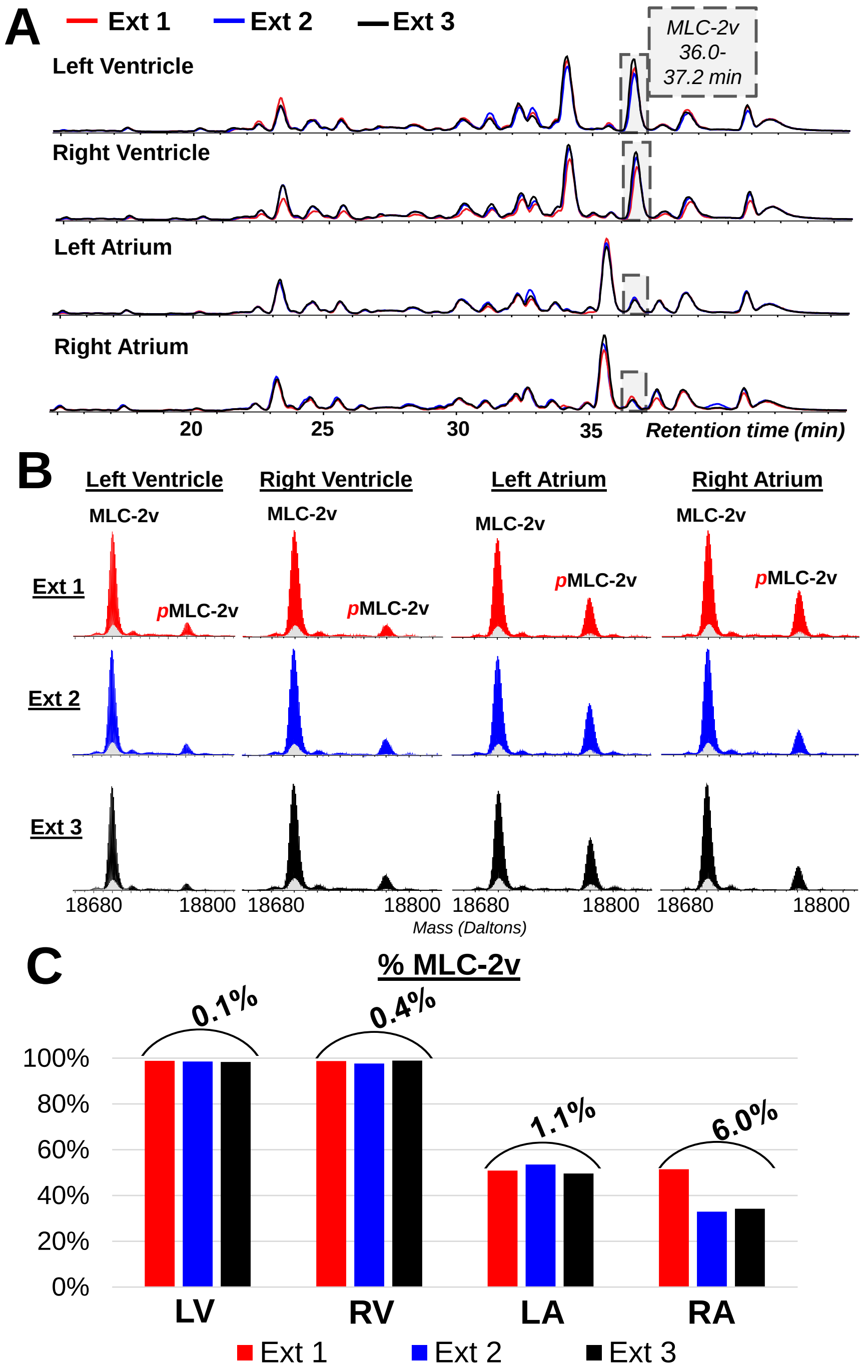


**Figure S5. Reproducible LC-MS analysis of sarcomeric proteins from multiple regions in the human heart. A)** BPCs are overlaid for (n=3) individual extraction replicates for LV, RV, LA, and RA. The retention time where ions corresponding to MLC-2v elute is highlighted with a grey box. **B)** Deconvoluted mass spectra of MLC-2v from extraction replicates. Mono-phosphorylated species is notated as *p*MLC-2v. **C)** Bar graph showing relative MLC-2v signal in each extraction replicate by AUC EICs. Relative Standard Deviation (% RSD) is reported for each measurement.

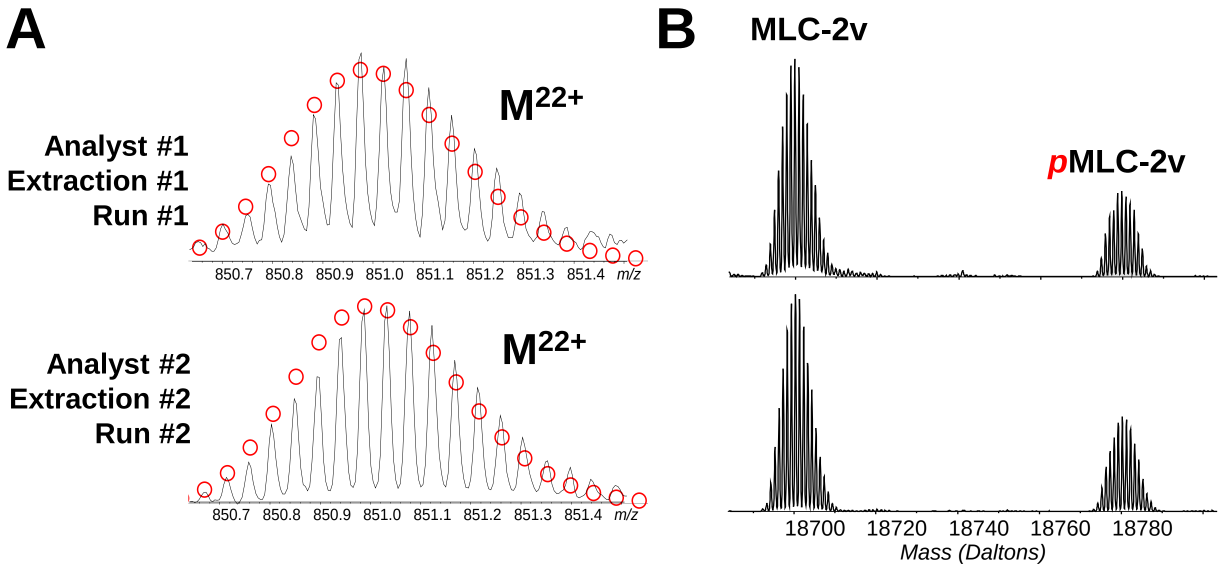


**Figure S6**. **MLC-2v detected in LA tissue across multiple extractions, sample runs, and analysts.** Extraction replicates of LA from the same donor heart were performed by different analysts, different buffers, and different LC-MS/MS runs. **A)** Zoomed in raw mass spectra of the most abundant charge state ($M^{22+}$) of MLC-2v shows good fitting of theoretical isotopic abundance, represented by red circles. **B)** Deconvoluted spectra of MLC-2v shows similar phosphorylation levels between measurements.

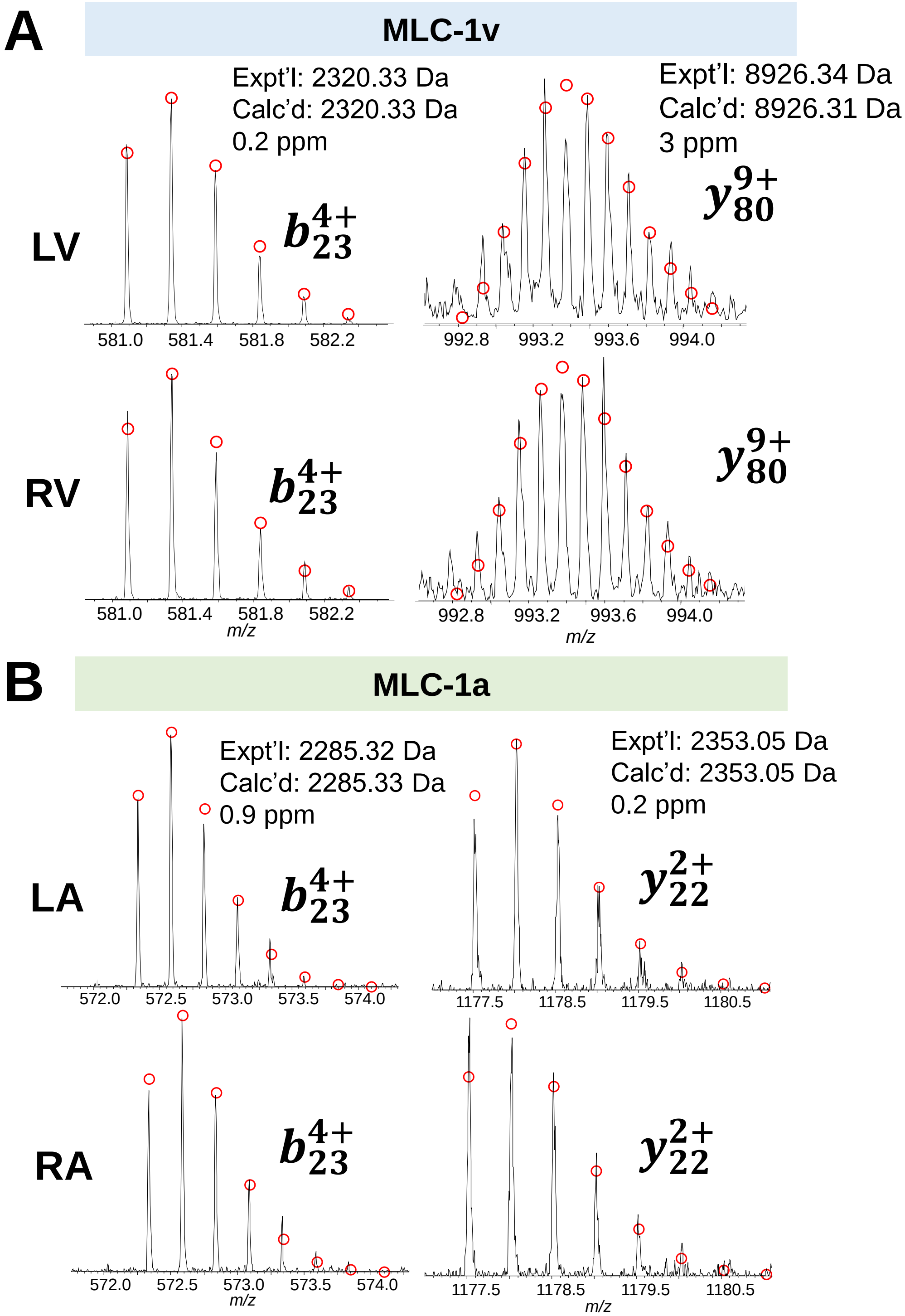


**Figure S7. MLC-1 Fragment Ions identified in ventricular and atrial tissue. A)** MLC-1v fragment ions were identified consistently in LV and RV generated by online CID. Theoretical isotopic distribution is demonstrated by red circles, and the protein experimental (Expt'l) and calculated (Calc'd) monoisotopic mass is reported. **B)** MLC-1a fragment ions were identified consistently across LA and RA.

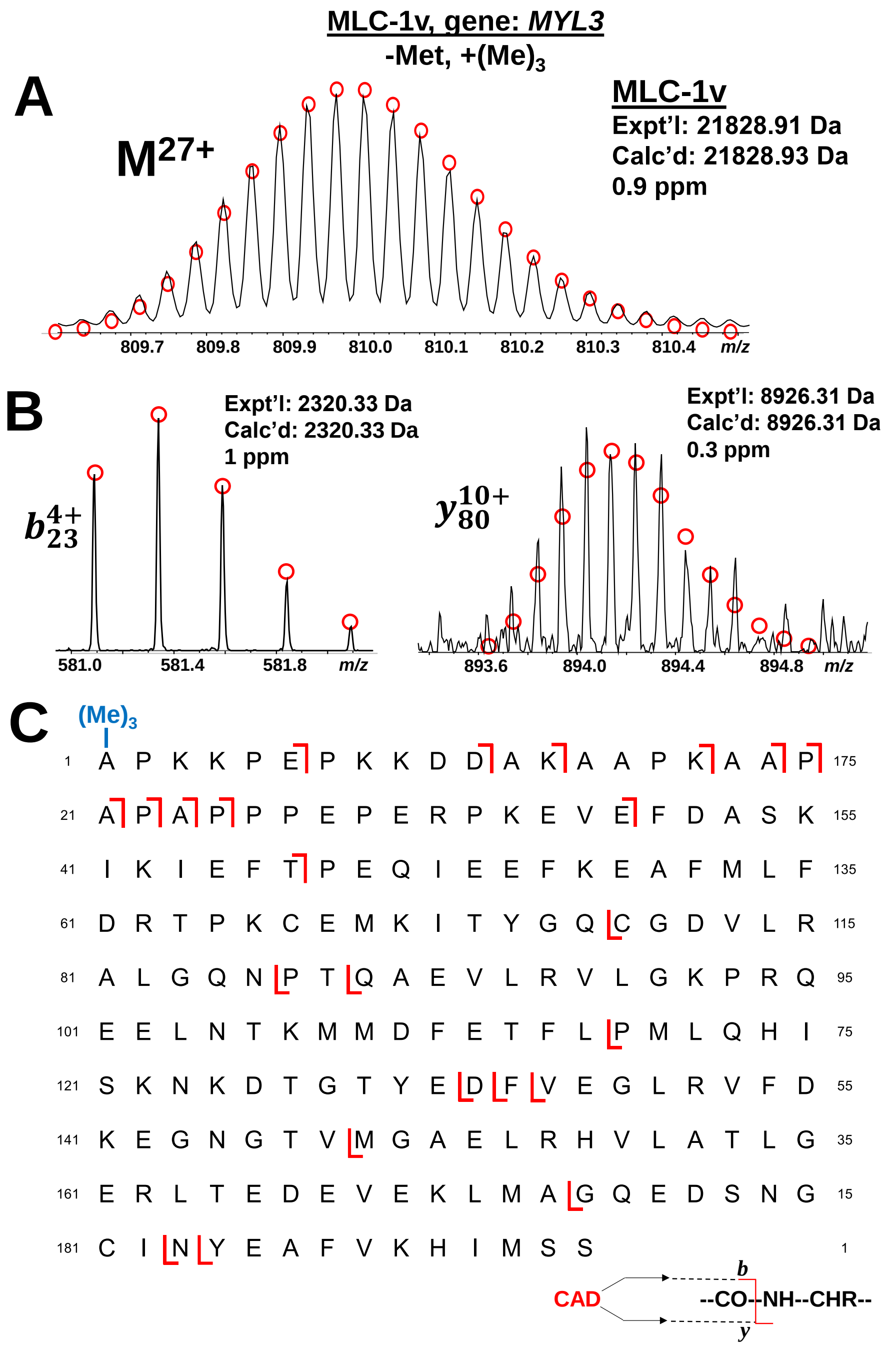


**Figure S8**. **Online LC-MS/MS of MLC-1v. A)** MLC-1v precursor ion shown at charge state $M^{27+}$. Theoretical isotopic distribution is demonstrated by red circles, and the protein experimental (Expt'l) and calculated (Calc'd) monoisotopic mass is reported. **B)** Representative fragment ion spectra produced by CID using 28.5 eV. N-terminal *b*23 fragment ion aligns with predicted mass accounting for N-terminal Met removal and trimethylation, and C-terminal *y*80 ion matches well with calculated mass without modification from canonical sequence. **C)** Sequence table showing bond cleavages of MLC-1v. Sequence accounts for the removal of N-terminal Methionine, and N-terminal trimethylation $(Me)_3$ is indicated in blue. 87% sequence coverage was achieved from a single LC-MS/MS run.

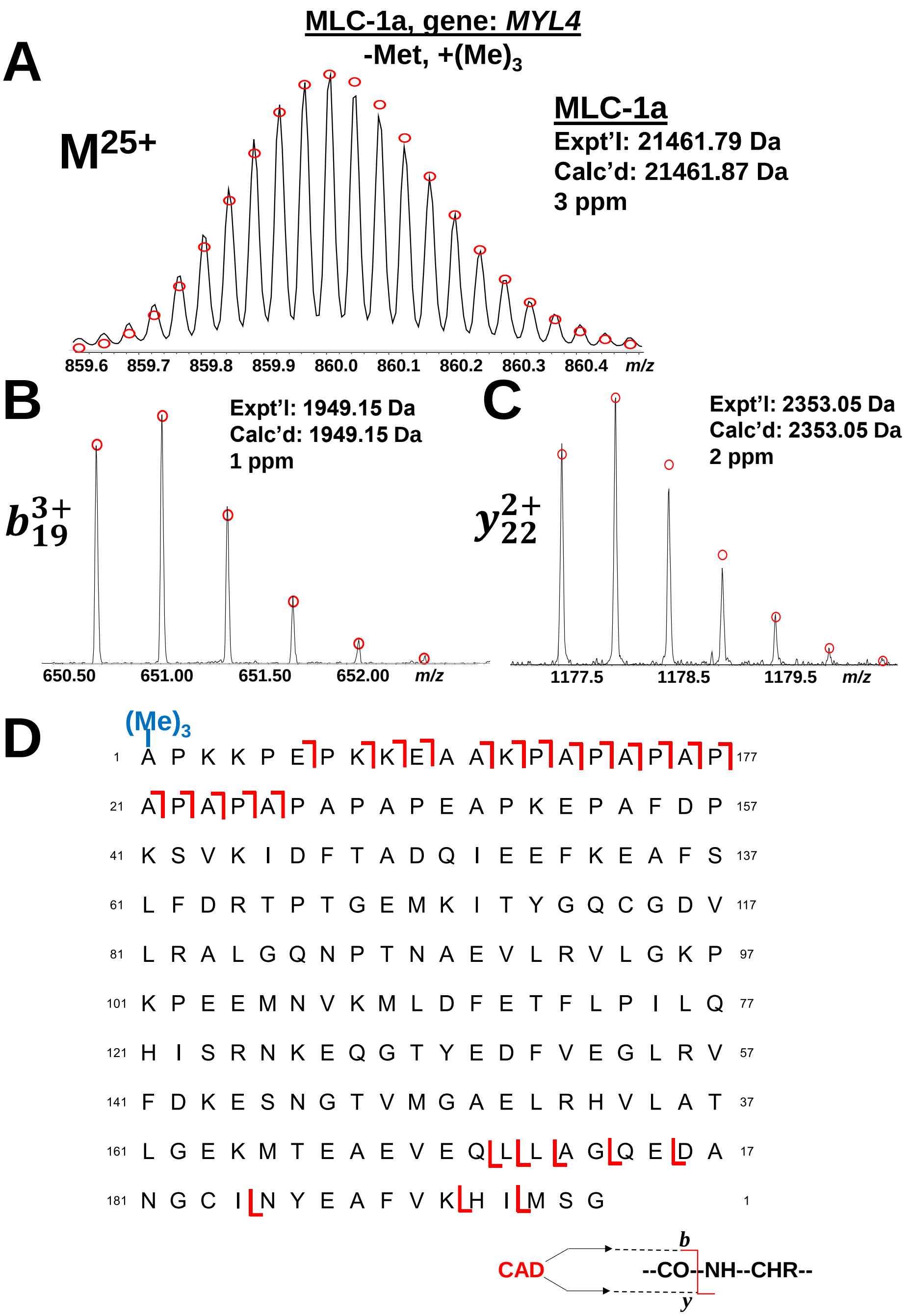


**Figure S9. Online LC-MS/MS of MLC-1a**. **A)** MLC-1a precursor ion shown at charge state 25+. Theoretical isotopic distribution is demonstrated by red circles, and calculated (Calc'd) and experimental (Expt'l) monoisotopic masses are given with error reported in parts per million (ppm). **B)** Representative fragment ion spectra produced by CID using 28.7 eV. N-terminal $b_{19}$ ion aligns with predicted mass accounting for N-terminal Met removal and trimethylation, and C-terminal $y_{22}$ matches well with calculated mass without modification from canonical sequence. **C)** Sequence table showing bond cleavages of MLC-1a. Sequence accounts for the removal of N-terminal Methionine, and N-terminal trimethylation $(Me)_3$ is indicated in blue. 26% sequence coverage was achieved through a single LC-MS/MS run.

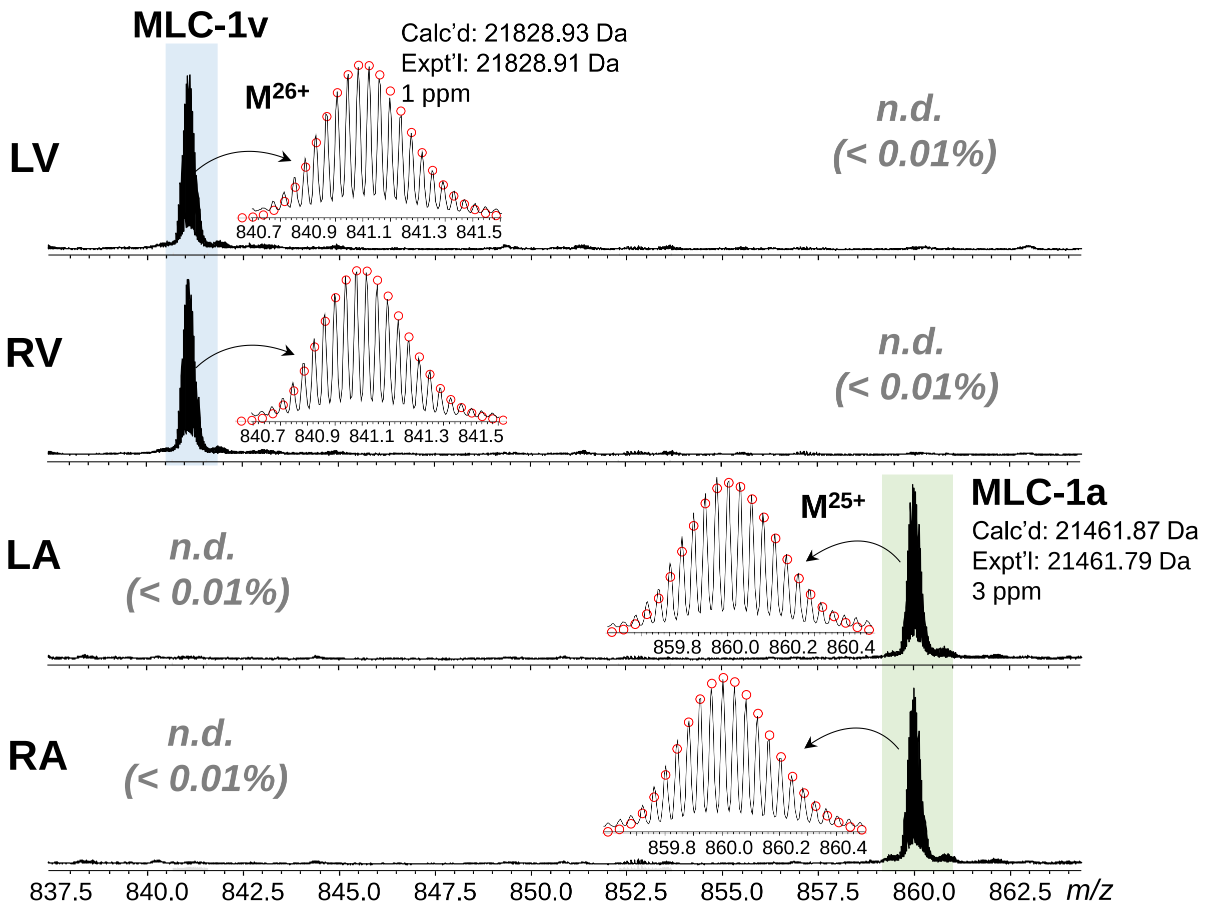


**Figure S10. Raw mass spectra of isoforms of MLC-1 in each of the four chambers of human donor hearts.** MLC-1v, shown at charge state $M^{26+}$ is the dominant isoform in LV and RV, and MLC-1a ($M^{25+}$) is the most abundant isoform in LA and RA. Inlays show zoomed mass spectra for selected charge states of MLC-1v and MLC-1a. Theoretical fitting of isotopic abundance and distribution of isotopomer peaks for each isoform is represented by red circles. Experimental (Expt'l) and calculated (Calc'd) monoisotopic masses with N-terminal Met removal and Trimethylation are reported (**Table S2**).

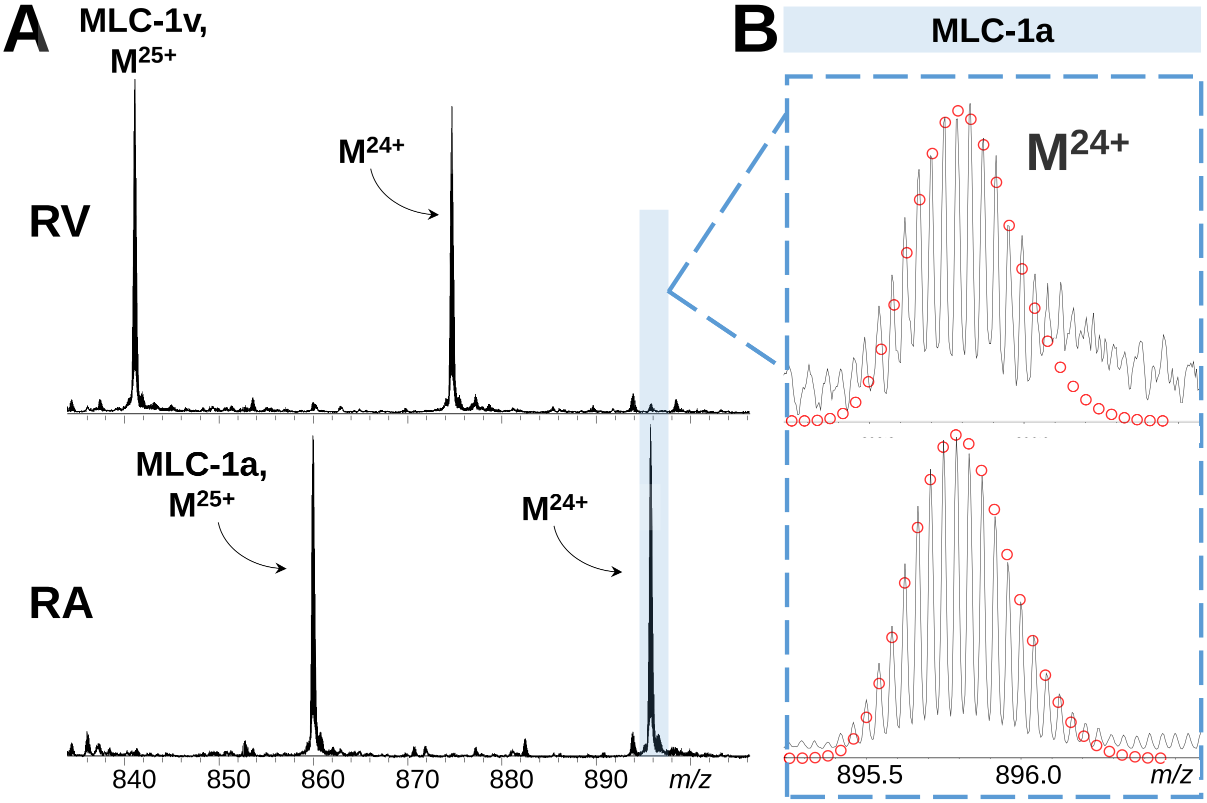


**Figure S11. MLC-1a detected at low levels in some ventricular donor tissues**. **A)** Mass spectra of MLC-1 isoforms in representative ventricular and atrial tissue. Charge state $M^{24+}$ for MLC-1a is highlighted in blue. **B)** Zoomed mass spectra showing the isotopic distribution and theoretical fit of charge state $M^{24+}$ of MLC-1a in the ventricular tissue (top) and atrial tissue (bottom).

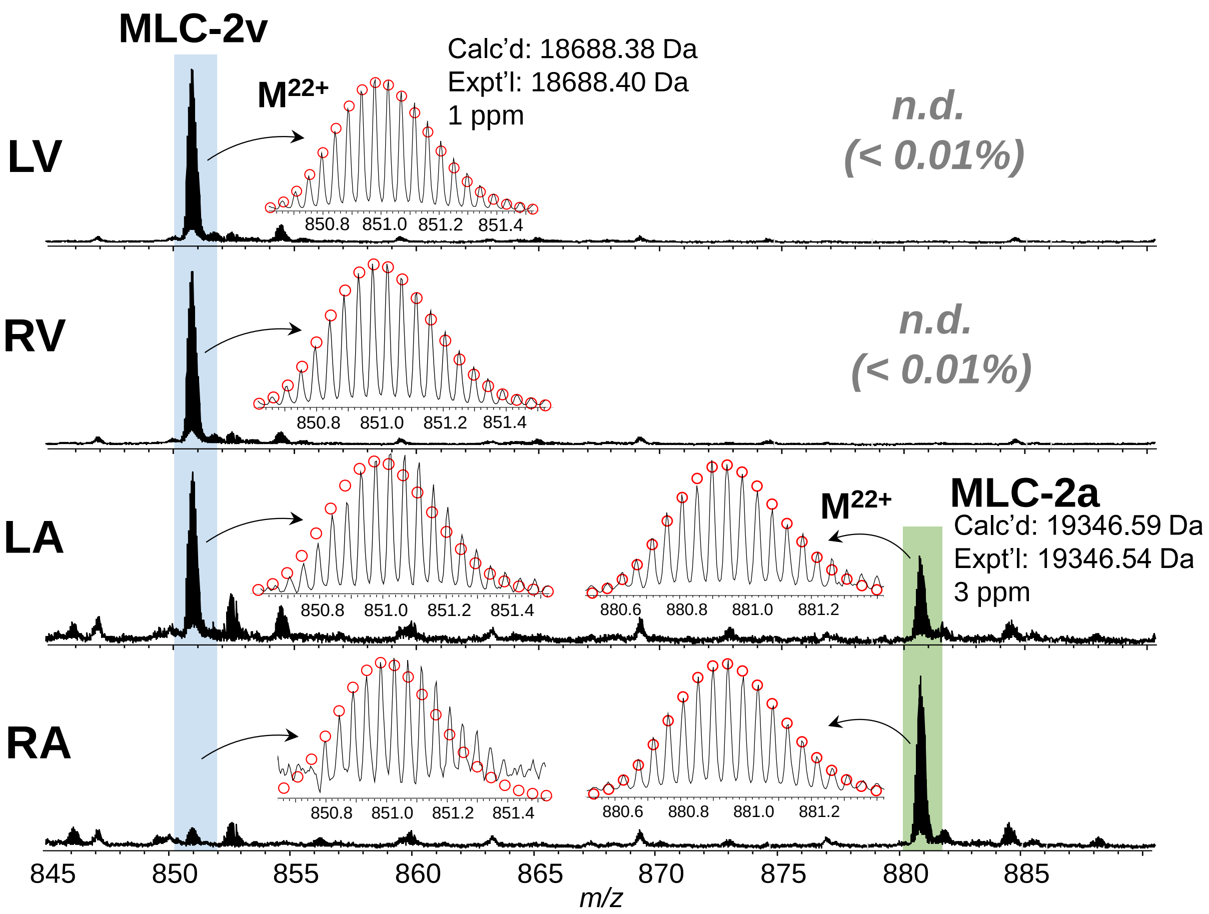


**Figure S12**. **Raw mass spectra of isoforms of MLC-2 in LV, RV, LA, and RA.** MLC-2v, shown at charge state $M^{22+}$ was detected across tissues from the four chambers of human donor hearts. MLC-2a ($M^{22+}$) is exclusively detected in LA and RA. Inlays show zoomed mass spectra for selected charge states of MLC-2v and MLC-2a. Theoretical fitting of isotopic abundance and distribution of isotopomer peaks is represented by red circles. Experimental (Expt'l) and calculated (Calc'd) monoisotopic masses are reported (**Table S2**).

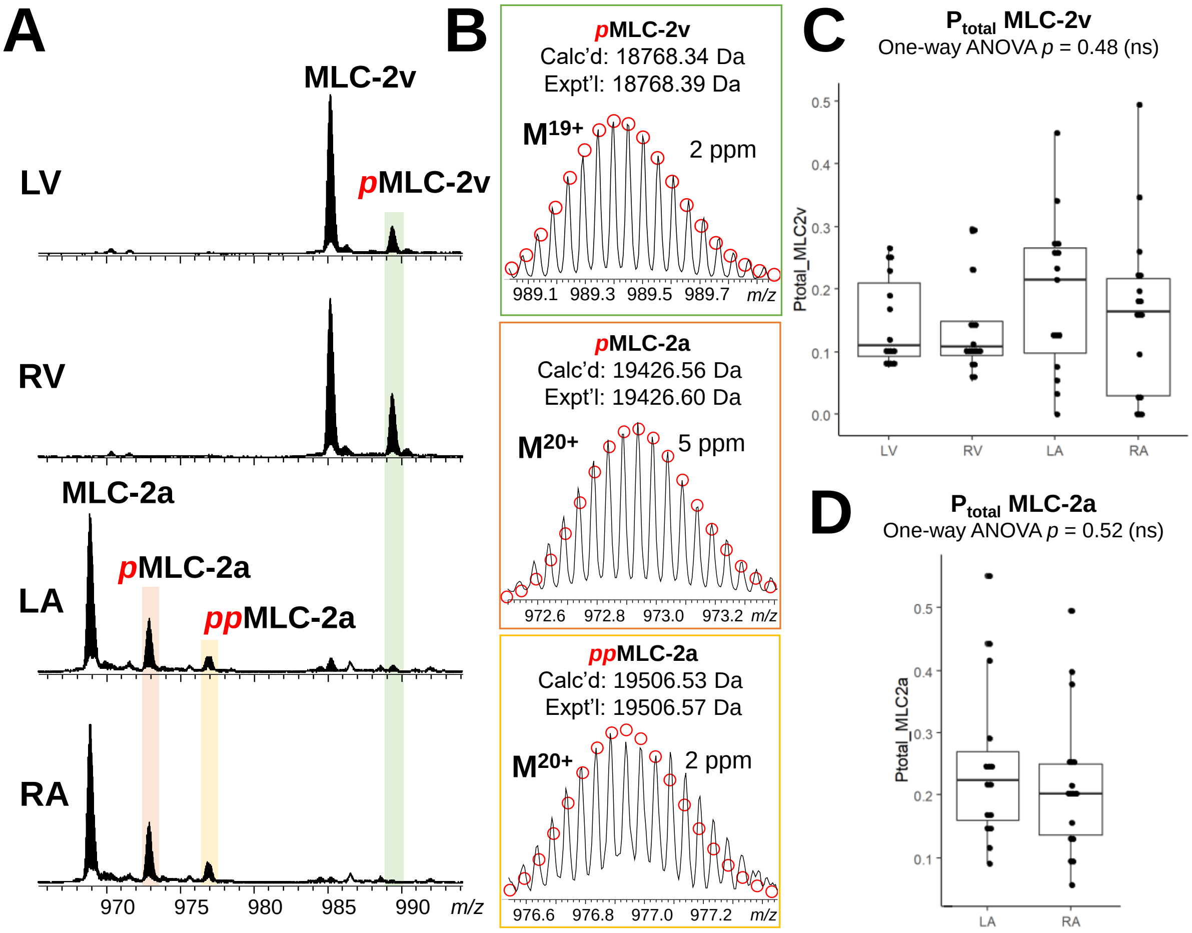


**Figure S13**. **Identification and quantitation of phosphorylated proteoforms of MLC-2 across the four cardiac chambers**. **A)** Raw mass spectra of MLC-2v ($M^{19+}$) and MLC-2a ($M^{20+}$) proteoforms in LV, RV, LA, and RA. Mono- and bis-phosphorylated proteoforms are notated by *p*MLC-2v, *p*MLC-2a, and *pp*MLC-2a, respectively. **B)** Zoomed mass spectra showing isotopic distribution of each phospho-proteoform are shown. Theoretical isotopic abundance of each proteoform is indicated by red circles and matched to experimental spectra. Experimental (Expt'l) and calculated (Calc'd) monoisotopic masses are reported for each proteoform (**Table S2**). **C)** Quantitation of relative phosphorylation of MLC-2 isoforms across the four chambers of the human heart. Relative phosphorylation, reported as $P_{total}$ values, was assessed by one-way ANOVA. No significant differences in relative phosphorylation for MLC-2v were detected ($p$ = 0.48) and **D)** MLC-2a ($p$ = 0.52).

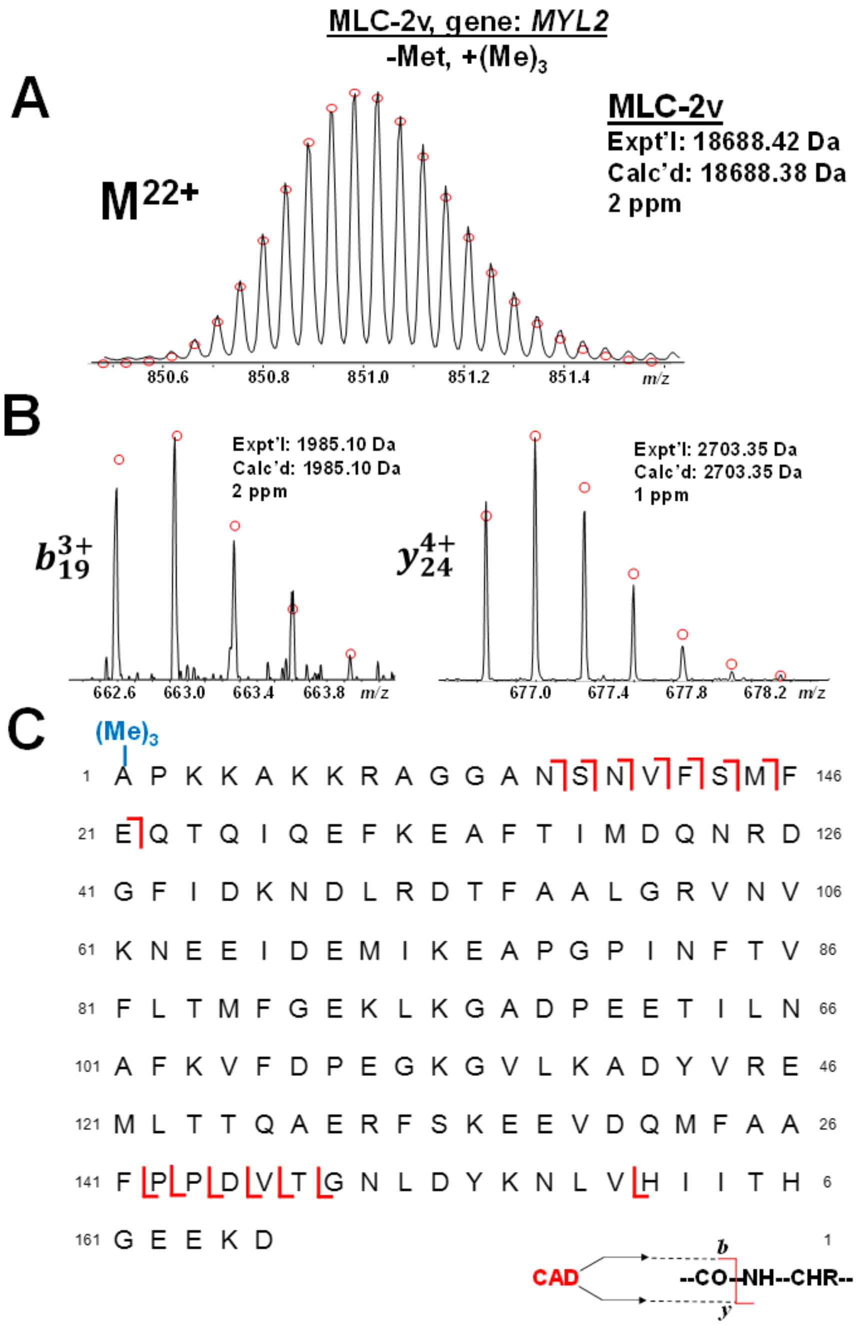


**Figure S14**. **Online LC-MS/MS of MLC-2v. A)** MLC-2v precursor ion shown at charge state $M^{27+}$. Theoretical isotopic distribution is demonstrated by red circles, and the protein experimental (Expt'l) and calculated (Calc'd) monoisotopic mass is reported. **B)** Representative fragment ion spectra produced by CID using 28.6 eV. N-terminal $b_{19}$ ion aligns with predicted mass accounting for N-terminal Met removal and trimethylation, and C-terminal $y_{24}$ ion matches well with calculated mass without modification from canonical sequence. **C)** Sequence table showing bond cleavages of MLC-2v. Sequence accounts for the removal of N-terminal Methionine, and N-terminal trimethylation $(Me)_3$ is indicated in blue. 31% sequence coverage was achieved from a single LC-MS/MS run.

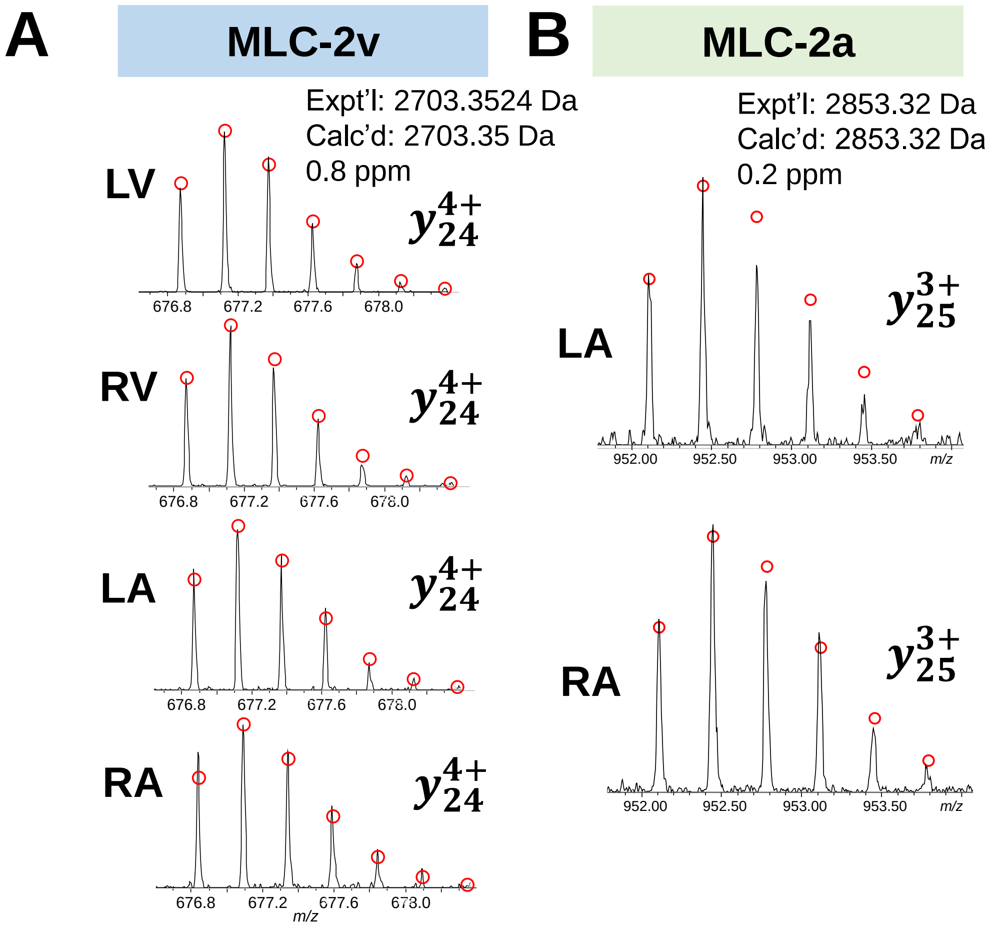


**Figure S15. MLC-2 fragment ions identified in ventricular and atrial tissue. A)** MLC-2v C-terminal fragment ions were identified consistently in LV and RV generated by online CID. Theoretical isotopic distribution is demonstrated by red circles, and the protein experimental (Expt'l) and calculated (Calc'd) monoisotopic mass is reported. **B)** MLC-2a fragment ions were identified consistently across LA and RA.

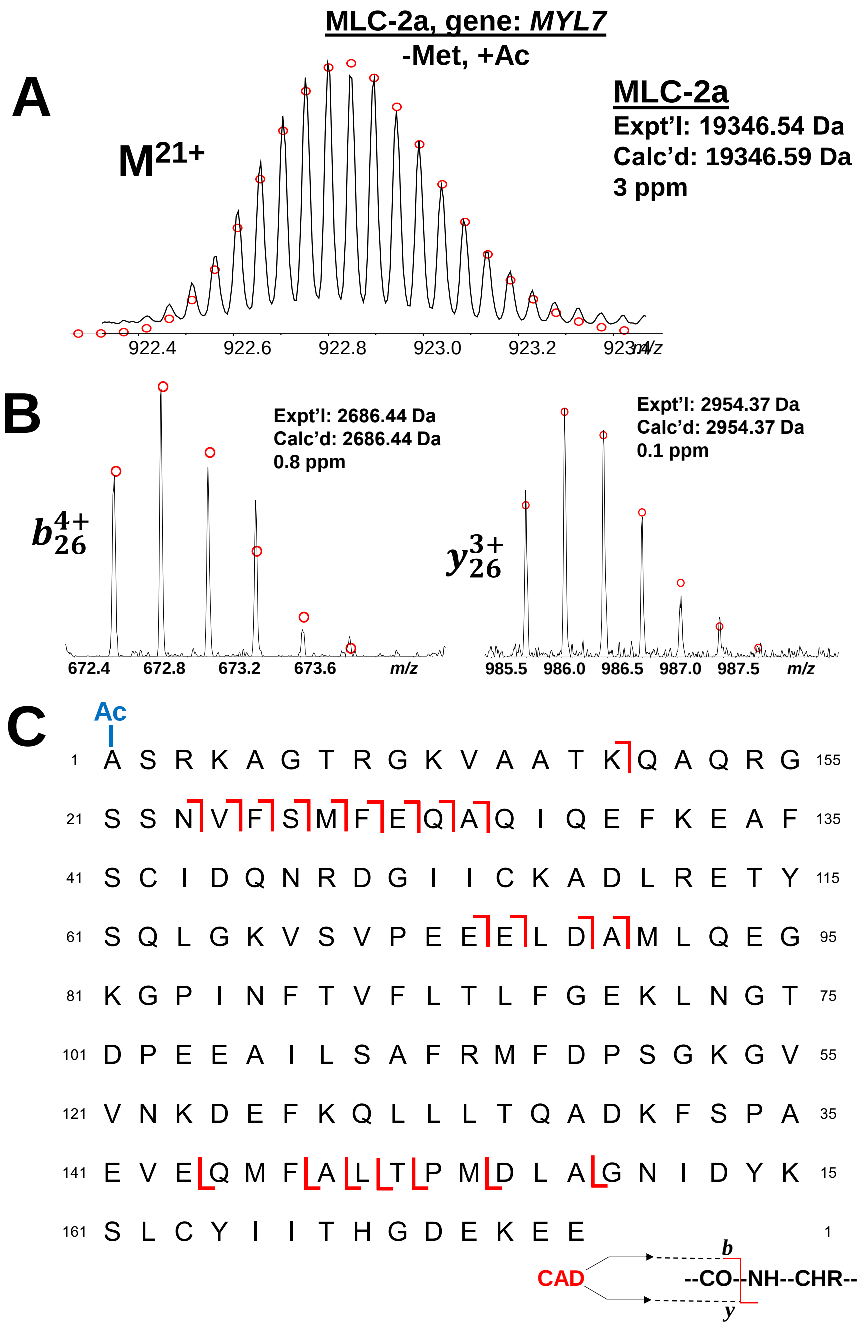


**Figure S16**. **Online LC-MS/MS of MLC-2a. A)** MLC-2a precursor ion shown at charge state $M^{21+}$. Theoretical isotopic distribution is demonstrated by red circles, and the protein experimental (Expt'l) and calculated (Calc'd) monoisotopic mass is reported. **B)** Representative fragment ion spectra produced by CID using 29 eV. N-terminal $b_{26}$ ion aligns with predicted mass accounting for N-terminal Met removal and Acetylation, and C-terminal $y_{26}$ ion matches well with calculated mass without modification from canonical sequence. **C)** Sequence table showing bond cleavages of MLC-2a. Sequence accounts for the removal of N-terminal Methionine, and N-terminal Acetylation (Ac) is indicated in blue. 61% sequence coverage was achieved from a single LC-MS/MS run.

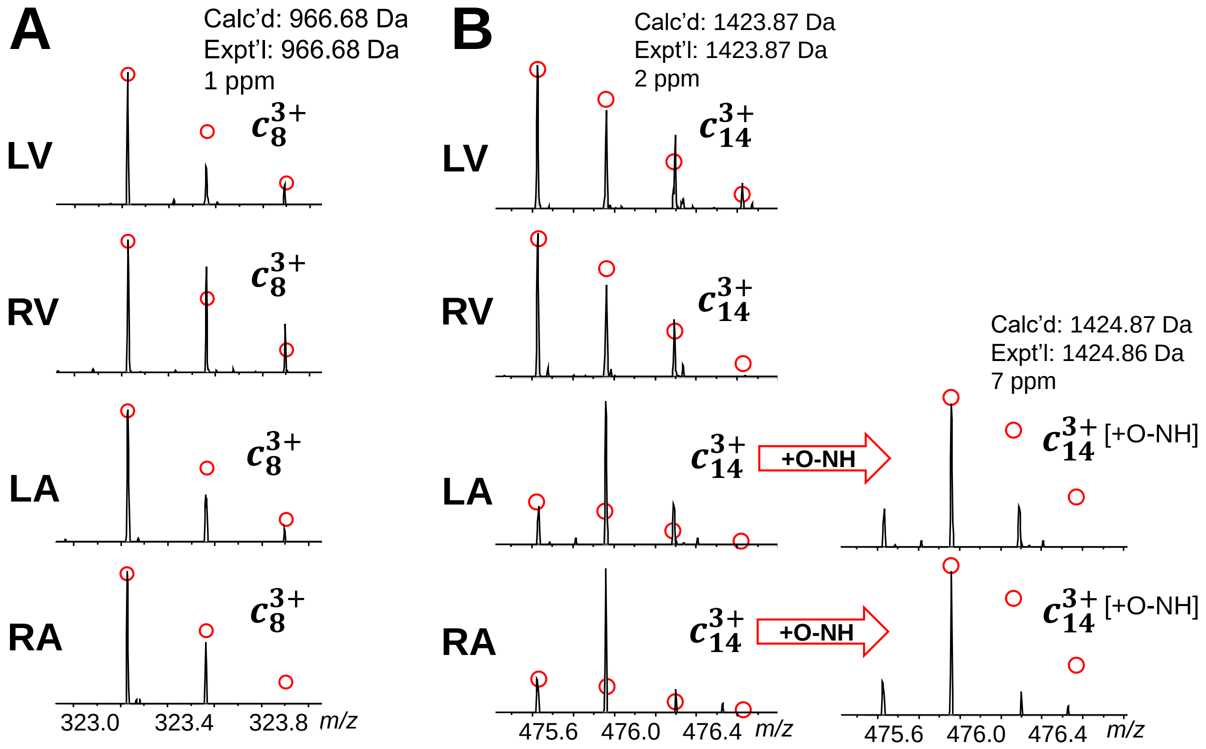


**Figure S17. MLC-2v fragment ions generated by Electron Transfer Dissociation (ETD) identified in ventricular and atrial tissue. A)** ETD fragmentation yielded smaller N-terminal fragments, and ion $c_8$ matches well with the predicted MLC-2v sequence and is consistent with observed fragments in LV and RV. **B)** Fragment ion $c_{14}$ showed a reduction of monoisotopic peak intensity in LA and RA similarly to the CID spectra. Fragment ions in LA and RA align closely with the alternative theoretical distrubtion accounting for deamidation.
